# Astroglial gap junctions strengthen hippocampal network activity by sustaining afterhyperpolarization via KCNQ channels

**DOI:** 10.1101/2022.12.14.520502

**Authors:** Elena Dossi, Lou Zonca, Helena Pivonkova, Lydia Vargova, Oana Chever, David Holcman, Nathalie Rouach

**Author notes:** Astrocytes and Vascular Niche, Marianne Grunberg-Manago Research Center, University of Rouen-Normandy, Mont-Saint-Aignan Campus, INSERM U1239, Rouen, France (Oana Chever). These authors contributed equally (Elena Dossi, Lou Zonca, David Holcman, Nathalie Rouach).

## Abstract

Throughout the brain, astrocytes form networks mediated by gap-junction channels that promote the activity of neuronal ensembles. Although their inputs on neuronal information processing are well established, how molecularly gap junction channels shape neuronal network patterns remains unclear. Here using astroglial connexin-deficient mice, in which astrocytes are disconnected and neuronal bursting patterns are abnormal, we found that astrocyte networks strengthen bursting activity via dynamic regulation of extracellular potassium levels, independently of glutamate homeostasis or metabolic support. Using a novel facilitation-depression model, we identified neuronal afterhyperpolarization as the key parameter underlying bursting patterns regulation by extracellular potassium in mice with disconnected astrocytes. We confirmed experimentally this prediction, and revealed that astroglial network-control of extracellular potassium sustains neuronal afterhyperpolarization via activation of KCNQ voltage-gated K^+^ channels. Altogether, these data delineate how astroglial gap-junctions mechanistically strengthen neuronal population bursts, and points to approaches for controlling aberrant activity in neurological diseases.

## Introduction

Astrocytes are instrumental components of the tripartite synapse playing essential roles in brain information processing. Primarily organized in intercellular networks via gap junctions (GJ), astrocytes communicate directly with neurons and other astrocytes to integrate neuronal activity via activation of their channels, receptors and transporters (Dallérac et al., 2013; Perea et al., 2009). Astrocytes can in turn modulate neuronal excitability, synaptic transmission and plasticity via several mechanisms, including uptake of ions and neurotransmitters, release of gliotransmitters, and physical coverage of synapses (Dallérac et al., 2013; Perea et al., 2009).

GJ are highly specialized channels formed by two main connexin (Cx) subunits, Cx43, which is expressed from embryonic to adult stages, and Cx30, which becomes expressed later in development (Giaume et al., 2010). They allow intercellular trafficking and long-range exchange of various ions, metabolites and neuromodulators up to 1.5 kD, and are important for redistributing energy metabolites to neurons and maintaining extracellular ion, neurotransmitter, and volume homeostasis (Amzica et al., 2002; Pannasch and Rouach, 2013). Thereby, GJ-connected astrocytic networks control basal synaptic transmission and plasticity (Pannasch et al., 2011; Rouach et al., 2008). Mice deficient in astroglial Cx43 and Cx30, hereafter referred to as Cx-deficient mice, have disconnected astrocytes, resulting in increased hippocampal synaptic transmission and reduced long-term synaptic plasticity, due to impaired astroglial glutamate and potassium clearance and regulation of extracellular space volume (Pannasch et al., 2011). Astrocytic networks also control neuronal population activity by promoting sustained coordinated bursts (Chever et al., 2016; Pannasch et al., 2012; Wallraff, 2006). Specifically, in Cx-deficient mice, the astrocytic disconnection results in shorter, but more frequent population bursts, which translate *in vivo* into a reduced severity of evoked seizures and associated convulsive behavior (Chever et al., 2016). This indicates that GJ-mediated astroglial networks can exacerbate pathological network activity and that a better understanding of the underlying mechanisms might inform therapeutic development of more effective anticonvulsive therapies. To date, the cellular and molecular mechanisms that underlie changes in bursting pattern in Cx-deficient mice remain poorly understood. In mice with disconnected astrocytes, the altered bursting pattern is associated with increased synaptic noise, leading to a depolarized neuronal resting membrane potential, a decreased neuronal release probability, and impaired synchronization (Chever et al., 2016). In addition, neuronal excitability is increased and the refractory period after bursting is decreased. Although these data provided insight into multiple neuronal and synaptic parameters impacted by Cx- deficiency, we lack an understanding of their relative contribution to the changed bursting pattern and of the underlying astroglial mechanism.

Thus, we here investigated the mechanisms by which astrocytic networks modulate neuronal network bursts dynamics. By combining experimental and modeling approaches, we found that astroglial networks regulate bursting patterns by controlling neuronal afterhyperpolarization (AHP) via extracellular potassium modulation of KCNQ channel currents.

## Results

### Astroglial gap junctions regulate extracellular potassium levels during bursting

GJ contribute to extracellular homeostasis of various ions, neurotransmitters and metabolites such as potassium ions (K^+^), glutamate and glucose (Dallérac et al., 2018; Pannasch and Rouach, 2013), respectively, but their physiological relevance is unclear. We here first investigated whether astroglial GJ contribute to alterations in extracellular K^+^ levels ([K^+^]_e_) during bursting activity. Astrocytes are densely populated with K^+^ channels. Their membrane potential reflects the presence of high resting conductances for K^+^ and is thus highly sensitive, with a quasi-nernstian relationship, to alterations in [K^+^]_e_ (Amzica, 2002; Amzica and Massimini, 2002). We therefore first assessed alterations in [K^+^]_e_ by recording astrocyte membrane potential using whole-cell patch clamp recordings. We observed that disconnected astrocytes in Cx-deficient mice displayed a depolarized resting membrane potential compared to wild type (WT) astrocytes (WT: -82.98 ± 0.37 mV, n = 17 cells from 4 mice, Cx-deficient: -79.67 ± 0.69 mV, n = 13 cells from 5 mice, p = 0.0001; Fig. 1A-B). We also found that during bursting activity, astrocyte membrane potential depolarizations displayed increased frequency in Cx-deficient relative to WT mice (/min: 5.81 ± 0.62 vs 3.16 ± 0.40, p = 0.0002), but were reduced in amplitude (ΔVm: 2.68 ± 0.35 mV vs 5.07 ± 0.86 mV, p = 0.0207) and duration (6.17 ± 0.91 s vs 9.41 ± 0.49 s, p = 0.0029, n =13 and 17 cells from 5 and 4 mice, respectively; Fig. 1A-B). Furthermore, astroglial membrane potential depolarizations evoked-synaptically by Schaffer collateral stimulation with increasing strength (0.1 and 0.5 ms pulse width) were also reduced in Cx-deficient relative to WT mice (area (mV*s): 0.1 ms: 7.79 ± 1.41 vs 20.76 ± 4.13, p = 0.0072, 0.5 ms: 10.65 ± 1.92 vs 24.25 ± 4.24, p = 0.0077; amplitude (mV): 0.1ms: 2.57 ± 0.31 vs 4.89 ± 0.82, p = 0.0359, 0.5 ms: 4.01 ± 0.46 vs 6.48 ± 0.83, p = 0.0406; n = 11 and n = 17 cells from 5 and 4 mice, respectively; Fig. 1C-D).

**Fig. 1.**
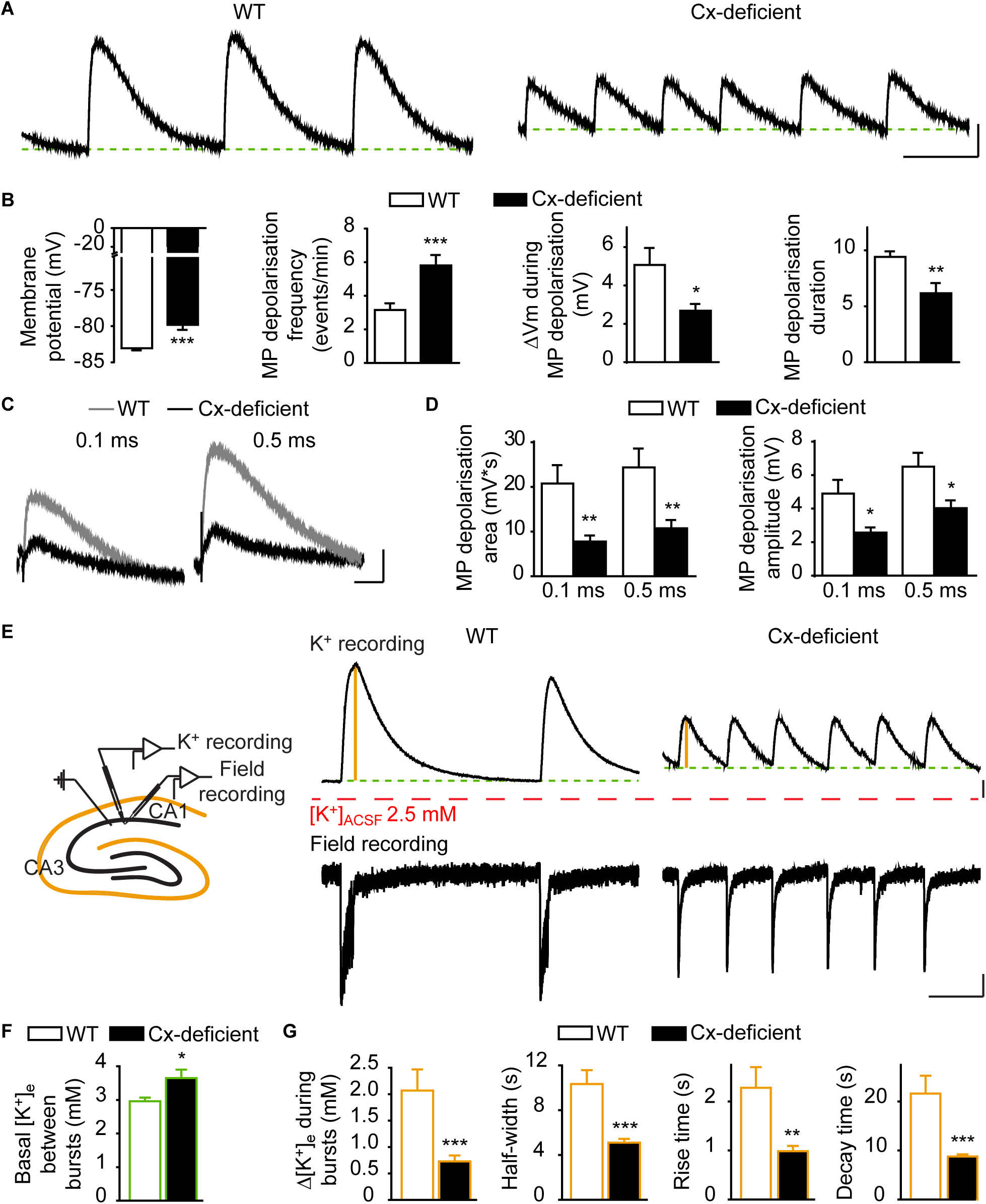
Gap junction-mediated astroglial networks modulate extracellular potassium levels during bursting. **A**) Representative traces of spontaneous astrocyte membrane depolarizations in WT (left) and Cx30^-/-^Cx43^fl/fl^ hGFAP-Cre (Cx-deficient, right) slices bathed in 0 Mg^2+^, picrotoxin ACSF and generating bursts. The green dashed lines indicate the resting membrane potential. Scale bars: 10 s, 2 mV. **B**) Quantification of CA1 *stratum radiatum* astrocyte resting membrane potential (MP), and spontaneous MP frequency, amplitude and duration in WT (17 cells from 4 mice) and Cx-deficient slices (WT: n = 17 cells form 4 mice; Cx-deficient: n = 13 cells from 5 mice; unpaired t-test for resting MP and spontaneous MP amplitude and duration, Mann-Whitney test for spontaneous MP frequency). **C**) Representative traces of evoked astrocyte response in WT (grey) and Cx-deficient (black) slices after 0.1 ms (left) and 0.5 ms (right) Schaffer collateral stimulation. Scale bars: 2 s, 2 mV. **D**) Quantification of MP depolarization area and amplitude in WT and Cx-deficient astrocytes after 0.1 and 0.5 ms Schaffer collateral stimulation (WT: n = 17 cells form 4 mice; Cx-deficient: n = 13 cells from 5 mice; unpaired t-test with Welch’s correction). **E**) Left, schematics illustrating simultaneous recordings of extracellular K^+^ and fEPSPs in the CA1 area of hippocampal slices. Right, representative traces from simultaneous recordings of K^+^ transients (top) and field potentials (bottom) in CA1 *stratum radiatum* of WT and Cx- deficient mice. The red and green dashed lines indicate K^+^ concentration in ACSF ([K^+^]_e_=2.5 mM) and basal extracellular K^+^ concentration measured between consecutive bursts, respectively. The orange bar indicates K^+^ changes (Δ[K^+^]_e_) recorded during bursts. Scale bars: 10 s, 0.2 mV (bottom)/0.5 mM (top). **F**) Quantification of basal [K^+^]_e_ between bursts, indicated by the green dashed line in panel e (WT: n = 16 slices from 9 mice; Cx-deficient: n = 17 slices from 6 mice; p < 0.05, unpaired t-test). **G**) Quantification of Δ[K^+^]_e_ during bursts (indicated by the orange line in panel e), half-width, rise time and decay time (WT: n = 13 slices from 8 mice; Cx-deficient: n = 18 slices from 9 mice; unpaired t-test with Welsch’s correction). Asterisks indicate statistical significance (*, p < 0.05; **, p < 0.01; ***, p < 0.001).

To directly assess alterations in [K^+^]_e_ during resting and bursting activity, we used K^+^- sensitive microelectrodes and electrophysiology to perform simultaneous recordings of [K^+^]_e_ and field potentials in Cx-deficient and WT hippocampal slices (Fig. 1E). We found that basal [K^+^]_e_ between bursts was higher in Cx-deficient relative to WT slices (3.65 ± 0.25 mM vs 2.96 ± 0.11 mM, p = 0.02 ; n = 16 and n = 17 slices from 9 WT and 6 Cx-deficient mice, respectively; Fig. 1F) and that burst-associated [K^+^]_e_ transients displayed reduced amplitude (0.73 ± 0.11 vs 2.07 ± 0.40 mM, p = 0.0009), half-width (5.09 ± 0.33 s vs 10.34 ± 1.23 s, p < 0.0001), rise time (0.99 ± 0.11 s vs 2.28 ± 0.42 s, p = 0.0019) and decay time (8.78 ± 0.43 s vs 21.63 ± 3.66 s, p = 0.0003; n = 18 and n = 13 slices from 9 and 8 mice, respectively; Fig. 1G), which reflected decreased burst strength in Cx-deficient mice (Fig. 1E). Altogether, these results indicate that disconnected astrocytes deficient for astroglial Cx impair extracellular K^+^ homeostasis, which results in increased basal [K^+^]_e_ and decreased [K^+^]_e_ rise during bursting activity.

### Manipulating extracellular K^+^ levels restores normal bursting pattern in Cx-deficient mice

We next investigated whether the perturbed extracellular K^+^ homeostasis results in the alteration of bursting activity found in Cx-deficient mice. To do so, we first tested in WT mice the impact of [K^+^]_e_ increase on bursting pattern. Using K^+^-sensitive microelectrodes, we found that basal [K^+^]_e_ increased by ∼ 0.7 mM between bursts in Cx-deficient slices (Fig. 1F), but these electrodes have limited access to the actual local [K^+^]_e_ in tissue compartments. Given that penetration and diffusion of exogenous K^+^ is limited in slices, extrapolating the actual local [K^+^]_e_ increase in Cx-deficient slices is challenging. Resting membrane potential of neurons is sensitive to [K^+^]_e_ and was previously reported to be depolarized by ∼ 5 mV in mice with disconnected astrocytes (Chever et al., 2016). We thus searched the exogenous [K^+^]_e_ needed to mimic this depolarization in WT slices, and found that increasing [K^+^]_e_ by 3.5 mM depolarized pyramidal cells by ∼ 5 mV (from - 62.7 ± 0.6 mV to - 57.6 ± 1.5 mV, n = 5; p < 0.001). Remarkably, the same increase in [K^+^]_e_ in WT slices strongly augmented burst frequency (from 2.28 ± 0.26 to 6.47 ± 0.43 bursts/min, n = 9 slices from 4 mice; p < 0.0001), and decreased burst duration (from 2.18 ± 0.22 s to 1.52 ± 0.07 s, n = 9 slices from 4 mice; p = 0.0065; Fig. 2A-B), thus mimicking changes induced by disconnection of astrocytes in Cx-deficient mice. Conversely, we then tested in Cx-deficient mice the impact of [K^+^]_e_ decrease on bursting pattern. We found that decreasing [K^+^]_e_ by 1 mM reduced burst frequency (from 5.67 ± 0.55 to 3.7 ± 0.32 bursts/min, n = 6 slices from 4 mice; p = 0.0018), as well as increased burst duration (from 1.23 ± 0.06 s to 1.76 ± 0.17 s, n = 6 slices from 4 mice; p = 0.0115; Fig. 2C-D), thus rescuing wild type bursting pattern. Altogether, our data show that relevant changes in basal [K^+^]_e_ can fully switch WT and Cx-deficient bursting pattern.

**Fig. 2.**
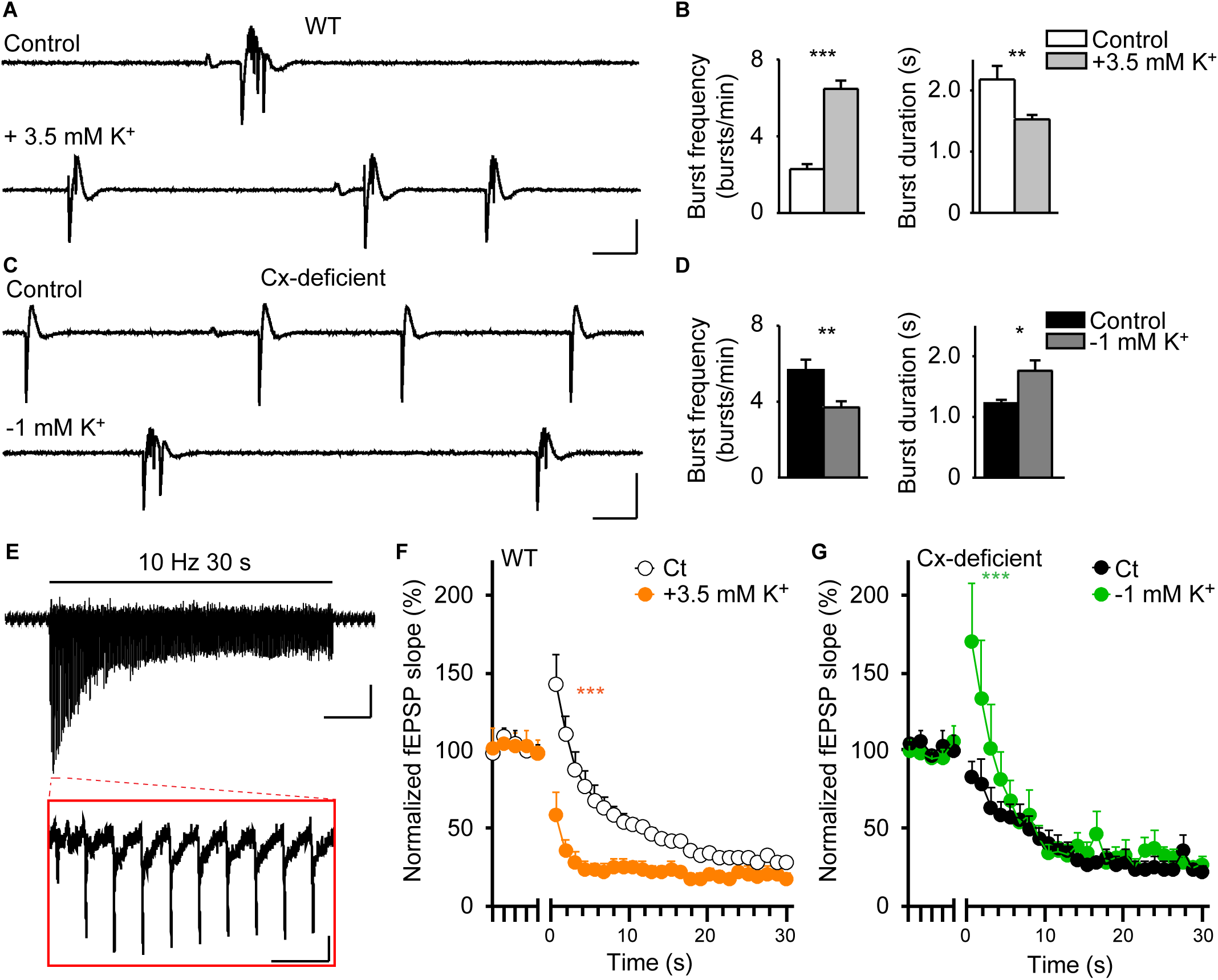
Alterations of extracellular potassium levels switch bursting patterns between wild type and astroglial Cx-deficient mice. **A**) Representative traces of hippocampal CA1 bursts recorded extracellularly in WT mice in control ACSF (top) and ACSF supplemented with 3.5 mM KCl (+ 3.5 mM K^+^, bottom). Scale bars: 2 s, 400 μV. **B**) Quantification of burst frequency and duration (n = 9 slices from 4 mice; paired t-test). **C**) Representative traces of hippocampal CA1 bursts in Cx-deficient mice in control ACSF (top) and in ACSF depleted by 1 mM KCl (-1 mM K^+^, bottom). Scale bars: 2 s, 400 μV. **D**) Quantification of burst frequency and duration (n = 6 slices from 4 mice; paired t-test). **E**) Top, Representative trace of fEPSPs evoked by repetitive stimulation (10 Hz, 30 s) of CA1 Schaffer collaterals. Scale bars: 5 s, 0.2 mV. Bottom, enlarged view of fEPSPs evoked by the first 10 stimuli. Scale bars: 200 ms, 0.2 mV. **F-G**) Quantification of relative changes in fEPSP slope induced by the 10 Hz stimulation over baseline responses measured before the onset of stimulation, in WT mice (**F**) in control ACSF (white) and ACSF supplemented with 3.5 mM KCl (orange; n = 9 slices; repeated measures two-way ANOVA), and in Cx-deficient mice (**G**) in control ACSF (black) and in ACSF depleted by 1 mM KCl (green; n = 7 slices; repeated measures two-way ANOVA). Asterisks indicate statistical significance (*, p < 0.05; **, p < 0.01; ***, p < 0.0001).

The altered bursting pattern in Cx-deficient mice is associated with a decreased neurotrasmitter release probability (Chever et al., 2016), which represents a hallmark of synaptic depression (Zucker and Regehr, 2002). Such release probability is correlated with the size of the vesicle readily releasable pool (Dobrunz and Stevens, 1997), determining in turn the recruitment of neurons during bursting activity (Cohen and Segal, 2011; Staley et al., 1998), which we also found to be reduced in Cx-deficient mice (Chever et al., 2016). To further test the implication of [K^+^]_e_ alterations in the change of bursting pattern in Cx-deficient mice, we here tested whether they also contribute to the synaptic depression. To do so, we performed repetitive stimulation of Schaffer collaterals (10 Hz, 30 s), which induces a rapid synaptic facilitation followed by a depression, resulting from presynaptic glutamate depletion (Fig. 2E). In Cx-deficient slices, we observed less facilitation and faster depression relative to WT slices (p < 0.0001 and p = 0.0436, n = 7 and 9 slices for Cx-deficient and WT mice, respectively; Fig. 2F-G). Remarkably, increasing [K^+^]_e_ by 3.5 mM in WT slices significantly reduced the initial facilitation (p < 0.0001) to Cx-deficient levels (p = 0.1630) and accelerated synaptic depression (p < 0.001; n = 9 slices; Fig. 2F, orange). Conversely, decreasing [K^+^]_e_ by 1 mM in Cx-deficient slices increased the initial facilitation and slowed down the subsequent depression, effectively mimicking the synaptic depression response in WT slices (p < 0.0001 and p = 0.0428, n = 7 slices; Fig. 2F-G). These data indicate that changes in basal [K^+^]_e_ can switch synaptic depression in WT and Cx-deficient mice.

### Impaired extracellular glutamate homeostasis or metabolic support does not mimick the alteration of bursting pattern in astroglial Cx-deficient mice

Astroglial networks also play a key role in restraining basal synaptic activity through modulation of extracellular glutamate clearance rate. Astroglial Cx-deficient mice indeed show increased excitatory synaptic activity due to an impairment of synaptic glutamate uptake, which makes glutamate persist longer at the synapse and prolong its activation of AMPA and NMDA receptors (Pannasch et al., 2011). We next asked if the altered bursting pattern in Cx-deficient mice results from persistent activation of glutamatergic receptors. To this end, we first prolongedglutamate receptor activation in WT slices by blocking AMPA receptor (AMPAR) desensitization with cyclothiazide (100 µM) (Fig. 3A) and found that it decreased burst frequency (from 2.78 ± 0.25 to 1.72 ± 0.27 bursts/min, p = 0.0022; n = 10 slices from 4 mice) with no effect on burst duration (control: 1.85 ± 0.13 s, cyclothiazide: 1.82 ± 0.11 s, p = 0.8816; n = 10 slices from 4 mice). Furthermore, and unlike in control condition, seizure-like events appeared after ∼20 min of cyclothiazide application with an average frequency of 0.37 ± 0.12 events/min (p = 0.0199; n = 10 slices from 4 mice) and duration of 12.39 ± 3.22 s (Fig. 3A-B). Thus, prolonged glutamate activation of AMPARs via inhibition of their desensitization did not reproduce the bursting pattern observed in slices from Cx-deficient mice. Conversely, decreasing glutamate activation of AMPARs in Cx-deficient slices by partial inhibition with NBQX at a low concentration (0.5 µM) decreased burst frequency (control: 5.99 ± 0.29 vs NBQX: 3.43 ± 0.38 bursts/min, p < 0.0001; n = 11 slices from 6 mice), but did not affect burst duration (control: 0.94 ± 0.04 s vs NBQX: 1.00 ± 0.04 s, p = 0.2882; n = 11 slices from 6 mice; Fig. 3C-D). Altogether, these results indicate that the impaired bursting pattern observed in Cx-deficient mice is unlikely resulting from the sole alteration in extracellular glutamate levels.

**Fig. 3.**
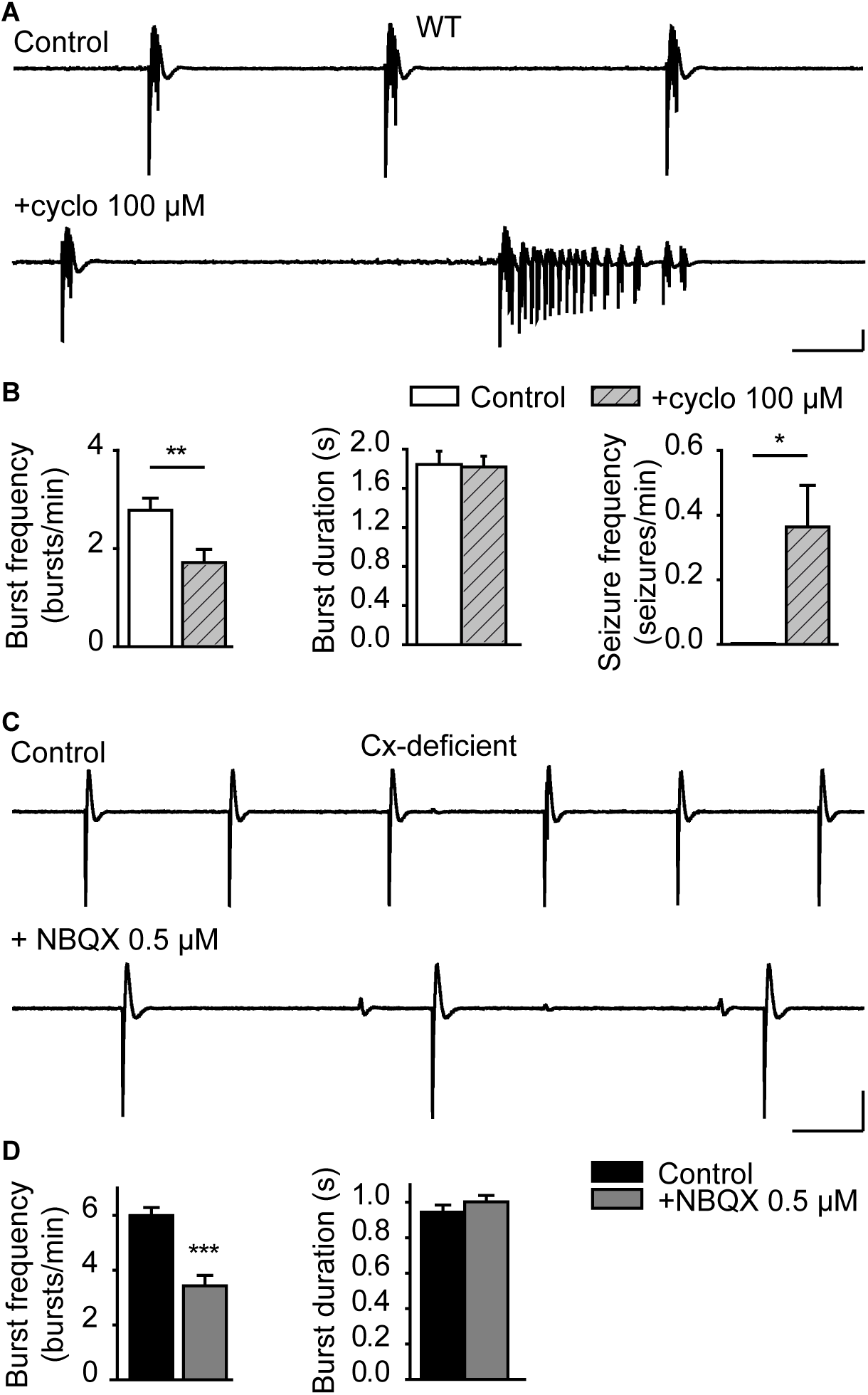
Impaired bursting in astroglial Cx-deficient mice is not due to altered extracellular glutamate homeostasis. **A**) Representative traces of hippocampal bursts in WT mice in control condition (top) and in presence of cyclothiazide (100 μM; bottom). Scale bars: 5 s, 200 μV. **B**) Quantification of burst frequency and duration and of seizure frequency (n = 10 slices from 4 mice; paired t-test). **C**) Representative traces of hippocampal bursts in Cx- deficient mice in control (top) and in presence of NBQX (0.5 μM; bottom). Scale bars: 5 s, 200 μV. **D**) Quantification of burst frequency and duration (n = 11 slices from 6 mice; paired t-test). Asterisks indicate statistical significance (*, p < 0.05; **, p < 0.01; ***, p < 0.0001).

Another key feature of astroglial networks is to provide energy metabolites to neurons via GJ-mediated intercellular pathway, thereby sustaining synaptic transmission and neuronal population bursts (Rouach et al., 2008). To test whether decreasing astroglial metabolite support contributes to the bursting pattern alteration in Cx-deficient mice, we deprived WT slices from glucose. We found that this treatment did not increase, but instead inhibited bursting activity, which displayed halved burst frequency after 20 min and was almost completely abolished after 30 min (control: 2.25 ± 0.31 vs 20 min: 1.24 ± 0.30 vs 30 min: 0.27 ± 0.07, p = 0.05 and p < 0.0001 for 20 and 30 min compared to Control; n = 11 slices; Fig. 4). This manipulation did not affect slice viability, as we could fully restore bursting activity by supplying exogenous glucose (11 mM, 20 min; 2.45 ± 0.40 bursts/min, p = 0.9290 compared to control; n = 11 slices; Fig. 4). Altogether, these data do not point to a role of impaired astroglial glucose supply in mediating the aberrant bursting pattern observed in Cx-deficient slices.

**Fig. 4.**
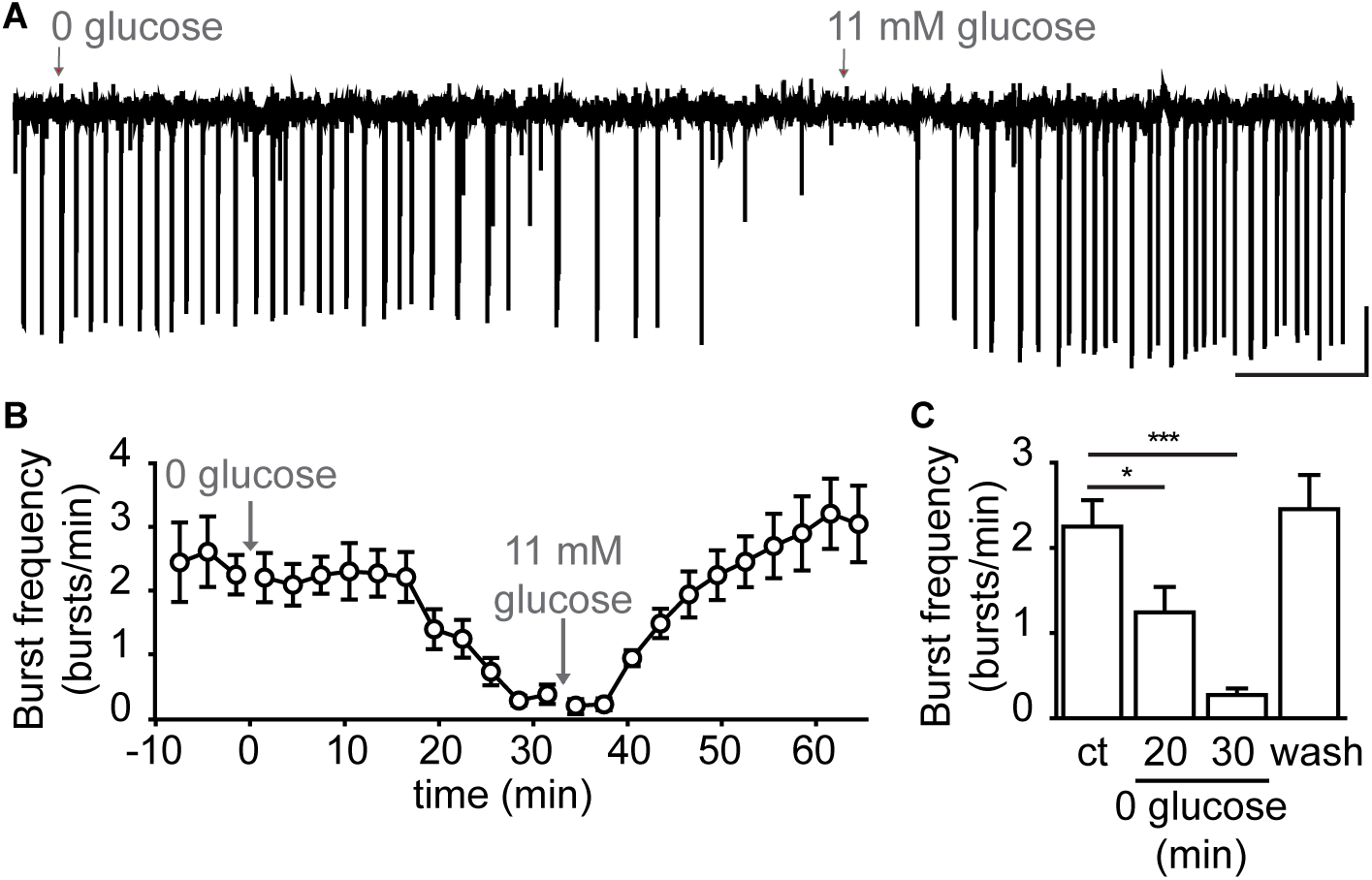
Impaired bursting in astroglial Cx-deficient mice is not due to altered metabolic support. **A**) Representative field potentialS recording of hippocampal bursting activity in WT mice during application of glucose-free ACSF (0 glucose) and washout in normal ACSF (11 mM glucose). Scale bars: 5 min, 0.2 mV. **B**) Temporal profile of burst frequency changes during perfusion of 0 glucose-ACSF (30 min) and washout in 11 mM glucose-containing ACSF (30 min) (n = 11 slices). **C**) Quantification of burst frequency in control, after 20 and 30 min in 0 glucose-ACSF and after 20 min of washout in normal ACSF (n = 11 slices; one- way ANOVA). Asterisks indicate statistical significance (*, p < 0.05; ***, p < 0.0001).

### Modeling neuronal bursting with astroglial networks

Our experimental data suggest that astroglial networks control bursting patterns via regulation of [K^+^]_e_ through an unknown mechanism. Extracellular K^+^ has multiple physiological targets, such as membrane potential and AHP, as well as synaptic noise and depression. In mice with disconnected astrocytes, all of these membrane and synaptic properties are altered ((Chever et al., 2016) and Fig. 2), but their relative contribution to the changes in bursting pattern is unknown. As there is no experimental tool to selectively target each of these components, we here used a modeling approach underlying burst generation. Based on experimental data from neuronal patch-clamp recordings of hippocampal pyramidal cells in WT and Cx-deficient mice, we developed a mean-field model that accounts for synaptic facilitation and depression and membrane properties, and in which we introduced an AHP component (Supplementary Fig. 1) (see Material and Methods). By using voltage time-series segmentation into three phases: bursting, AHP and quiescent phase (QP) (Supplementary Figs. 1A and 3A), we analyzed these electrophysiological data to generate associated duration histograms (Supplementary Fig. 1B). In Cx-deficient mice, we found that the distributions of burst durations and interburst intervals (IBI, which includes the AHP and QP) were shifted to lower values (burst durations: WT 3.06 ± 0.09 s, n = 284 bursts from 10 cells; Cx-deficient, 1.55 ± 0.02 s, n = 250 bursts from 6 cells; p < 0.0001; IBI: WT 18.02 ± 0.61 s, n = 284 bursts from 10 cells; Cx-deficient, 14.66 ± 0.31 s, n = 250 bursts from 6 cells; p = 0.0033; Supplementary Fig. 1B), and were also more peaked, reflecting reduced variance (burst duration: s_wt_ = 1.52 s, s_Cx-deficient_ = 0.37 s; IBI: s_wt_ = 10.27 s, s_Cx-deficient_ = 4.95 s). The AHP duration was also decreased in Cx-deficient mice (WT: 12.52 ± 0.56 s vs Cx-deficient: 8.03 ± 0.19 s, p < 0.0001, Supplementary Fig. 1B), while the QP duration was only slightly changed (WT: 5.50 ± 0.34 s vs Cx-deficient: 6.63 ± 0.32 s, p = 0.017; Supplementary Fig. 1B). Further, we found an increase in the synaptic noise amplitude (WT ² = 1.97 vs Cx-deficient ² = 3.80, Supplementary Fig. 1C), by computing the power spectral densities (PSD) of the QP using a Lorentzian fit on the low-pass filtered PSD at 30 Hz, as well as a depolarization of the membrane potential in Cx-deficient mice (WT: - 61.84 ± 1.43 mV, n = 12 cells; Cx-deficient, - 57.04 ± 0.56 mV, n = 10 cells; p = 0.0075; Supplementary Fig. 1D). To explore conditions under which modeling can reproduce experimental bursting data, we introduced a coarse-grained mathematical model that represents the dynamics of an averaged neuronal population driven by synaptic short-term plasticity. The model has three variables: the mean population membrane potential h, the short-term synaptic facilitation x and depression y variables.

The model can recapitulate four phases associated with a phenomenological burst decomposition (Fig. 5A):

1) Burst initiation (blue), dominated with fast dynamics driven by neuronal spiking and ending when the depression variable y reaches its minimum. The depression variable y can be interpreted as the change over time of the fraction of readily releasable pool of vesicles. Thus, at the minimum of y, the pool is depleted, and the bursting activity cannot be sustained anymore.
2) The recovery from the bursting phase (red, mid-burst) induces a membrane hyperpolarization and is dominated by the M-currents characterized by slower dynamics compared to the burst initiation. This phase lasts until the membrane hyperpolarization reaches a maximum value and the depression variable reaches an empirical threshold Y_h_.
3) The slow recovery of the membrane voltage to the resting potential (pink, burst refractory period): this phase is dominated by the slowest time scale and lasts until the voltage comes back around its resting value.
4) Membrane voltage fluctuations around the resting potential: this phase persists until the next burst starts (green, quiescent phase).

**Fig. 5.**
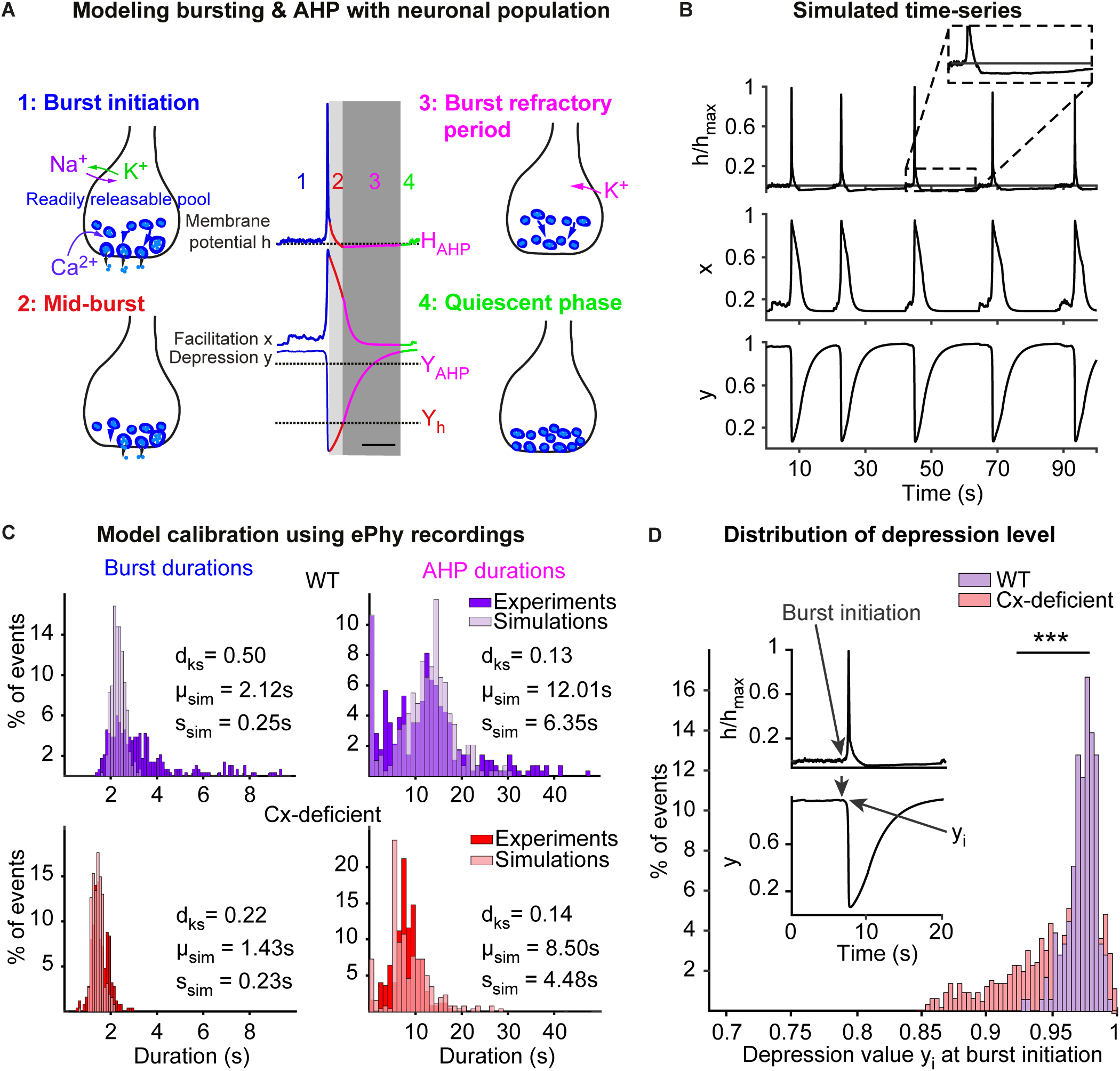
Modeling neuronal network bursting using depression-facilitation and AHP dynamics. **A**) Schematic decomposition of a burst, followed by AHP, into four steps: 1- Burst initiation mediated by facilitation and synaptic depression; 2- Recovery of synaptic depression until the maximum amplitude of hyperpolarization; 3- Burst refractory period. 4- Quiescent phase characterized by the lack of bursting activity. Scale bar: 5 s. **B**) Simulated time-series of normalized voltage (upper), facilitation x (center) and synaptic depression y (lower) over 100 s. **C**) Model calibration based on fitting simultaneously the experimental distributions of burst and AHP durations extracted from electrophysiological recordings segmentation (Supplementary Fig. 3a) of pyramidal neurons in WT (purple) and Cx-deficient (red) mice overlaid with the simulated distributions (light purple and red). **D**) Distribution of depression level y_i_ at burst initiation (arrow in the inset) obtained from 10000 s simulations for WT (purple) and Cx-deficient (red). Asterisks indicate statistical significance in two-sample Kolmogorov-Smirnov test (***, p < 0.0001). Parameters are summarized in Table 1.

**Table 1.** List and values of the model parameters.

|  | Parameters | Values |
| --- | --- | --- |
| $\sigma$ | Noise amplitude | 6 |
| $\tau$ | Fast time constant for $h$ | 0.05 s [this study] |
| $\tau_{m\text{AHP}}$ | Medium time constant for $h$ | 0.35 s (WT) / 0.15 s (Cx-deficient) [this study] |
| $\tau_{s\text{AHP}}$ | Slow time constant for $h$ | 10.5 s (WT) / 5 s (Cx-deficient) [this study] |
| $J$ | Synaptic connectivity | 4.21 [this study] |
| $K$ | Facilitation rate | 0.037 Hz modified: 0.04Hz in (Dao Duc et al., 2015) |
| $X$ | Facilitation resting value | 0.08825 modified: 0.05-0.1 in (Barak and Tsodyks, 2007b) |
| $L$ | Depression rate | 0.028 Hz modified: 0.037Hz in (Dao Duc et al., 2015) |
| $\tau_f$ | Facilitation time rate | 2.9 s modified 2-20s in (Dao Duc et al., 2015) |
| $\tau_r$ | Depression time rate | 0.9 s modified 1.3 in (Dao Duc et al., 2015) |
| $T_{\text{AHP}}$ | Undershoot threshold | - 30 (WT) / - 23 (Cx-deficient) [this study] |

Using this model, we simulated the mean membrane potential, the facilitation and the depression variables, and obtained time-series showing spontaneous bursts followed by AHP periods (Fig. 5B). The present model has 14 parameters, 6 of them are adapted from previous studies (Barak and Tsodyks, 2007a; Dao Duc et al., 2015; Holcman and Tsodyks, 2006; Tsodyks and Markram, 1997) and to fit the eight remaining ones, we calibrated the model by fitting optimally the distributions of burst and AHP durations both in the WT and in the Cx-deficient mice (Supplementary Fig. 2). If each distribution allows to constraint two parameters, then fitting the four histograms of burst and AHP durations in WT and Cx-deficient mice would give us unequivocally the 4x2=8 parameters that we need.

More specifically, we generated numerical simulations of equation (1) that we segmented using a thresholding method (Supplementary Fig. 3 and Supplementary Methods). We then obtained the statistics of bursting and AHP durations, which we fitted to generate distributions comparable to the ones extracted from segmenting the patch-clamp recordings (Supplementary Fig. 1A; see also Supplementary Fig. 3A for more details). To determine the optimal values of the parameters (Table 1), we first calibrated the model for the WT case (Fig. 5C, upper). We then modified the AHP duration, noise amplitude and depolarization by changing the medium τ*_mAHP_* and slow τ*_sAHP_* time constants, the AHP depth *T_AHP_*, the noise amplitude *σ* and the membrane depolarization *T*, in order to match the experimental distributions of the Cx-deficient case (Fig. 5C, lower). Finally, we did not use the distribution of the experimental QP durations but rather focused on fitting AHP durations (Supplementary Fig. 1B, lower panels), which carries most of the changes between WT and Cx-deficient mice.

After fitting, the model accounts for the shift in the distribution of burst durations from longer bursts in WT vs Cx-deficient mice (WT: 2.12 ± 0.25 s vs Cx-deficient: 1.43 ± 0.23 s). However, we could not account for the large variance of the distribution observed in the WT experimental data (Fig. 6C upper left, dark purple histogram). This limitation is probably due to the mean-field approximation (population level).

**Fig. 6.**
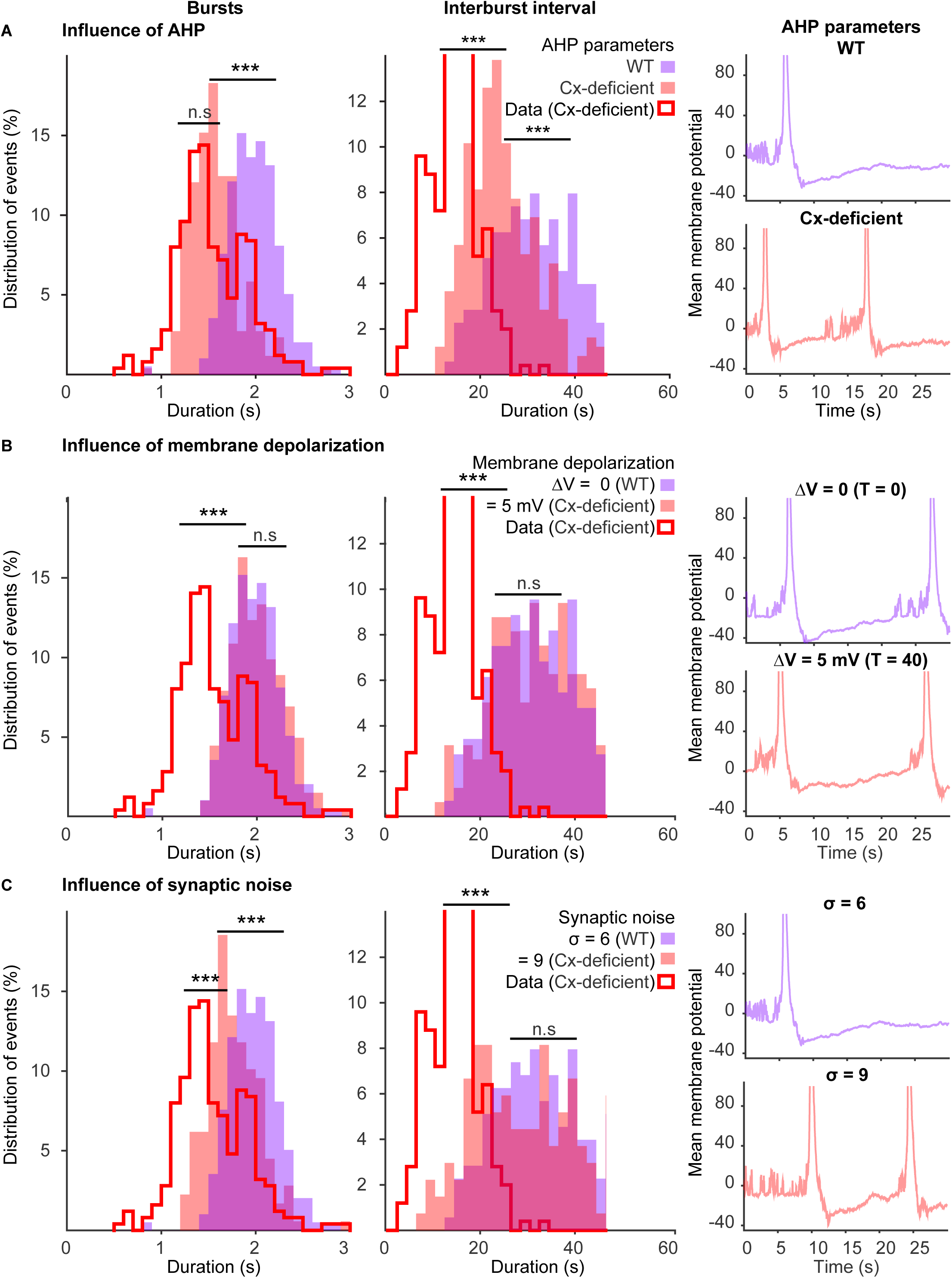
Relative contribution of AHP, synaptic noise and membrane potential depolarization on bursting dynamics. **A**) Left: distribution of burst (left) and IBI (right) durations for τ*_sAHP_* = 10.5 s, τ*_mAHP_* = 0.35 s and *T_AHP_* = - 30 (WT, light purple), and for τ*_sAHP_* = 5 s, τ*_mAHP_* _=_ 0.15s and *T_AHP_* = - 23 (Cx-deficient, light red). *σ* = 6 and *T* = 0 for both WT and Cx-deficient (5000 s simulations; p < 0.0001 for burst and IBI, Two-sample Kolmogorov-Smirnov test). The red curves represent the burst and IBI duration distributions obtained in the experimental data from Cx-deficient mice. Right: simulated mean voltage time series with AHP parameters of WT (top) and Cx-deficient (bottom) conditions. **B**) Same plots as panel **A** with τ*_sAHP_* = 10.5 s, τ*_mAHP_* = 0.35 s, *T_AHP_* = -30, *σ* = 6 and *T* = 0 (WT, light purple) or *T* = 40 (Cx-deficient, light red) (p = 0.88 for burst and p = 0.38 for IBI, Two-sample t- test). **c**) Same plots as panel **a** with τ*_sAHP_* = 10.5s, τ*_mAHP_* = 0.35s, *T_AHP_* = - 30, *T* = 0 and *σ* = 6 (WT, light purple) or *σ* = 9 (Cx-deficient, light red) (p < 0.0001 for bursts and p = 0.04 for IBI, Two-sample t- test). Asterisks indicate statistical significance (*, p < 0.05; ***, p <0.0001).

Further, since the experimental data indicated presence of synaptic depression in Cx-deficient mice (Fig. 2E-G), we investigated the depression level using the calibrated model. For that purpose, in the simulations, we reported the synaptic depression level y_i_ at the initiation of each detected burst (Fig. 5D, inset). With this approach, we found that neurons in astrocyte disconnection conditions are more depressed compared to control condition as the distribution of depression values is shifted to the left (p < 0.0001; Fig. 5D), in agreement with the experimental data. Altogether, our calibrated mean-field model accounts for the burst properties observed experimentally in WT and Cx-deficient mice.

### Modeling predicts that astroglial gap junctions set bursting patterns by regulating neuronal afterhyperpolarization

We next determined the relative contribution of the changes in either AHP, basal membrane depolarization, synaptic noise or depression to the Cx-deficient bursting pattern. To this end, we ran bursting simulations with parameters obtained in the fit of the WT distributions, except for the tested factor (AHP, noise amplitude or depolarization threshold), for which we used the data from the fit of the Cx-deficient condition (Fig. 6, red). These simulations were then compared to the ones computed using all parameters from the fit of WT distributions (Fig. 6, purple).

We first tested the selective effect of decreasing the AHP (medium τ_mAHP_ and slow τ_sAHP_ timescales and AHP depth T_AHP_), and found that durations of bursts and IBI shifted towards reduced values (p < 0.0001; Fig. 6A), mimicking the burst duration value we observed experimentally in the Cx-deficient condition (p = 0.23; Fig. 6A). Next, we investigated the selective impact of the membrane depolarization (ΔV = 5 mV), but found no shift in the distribution histograms for bursts and IBI durations (p = 0.78 and p = 0.60, respectively, Fig. 6B), thus resulting in a different pattern compared to the Cx-deficient condition (p < 0.0001, Fig. 6B). We also tested the specific effect of increasing the synaptic noise (σ, noise amplitude), and found that this only caused a shift in burst durations with no effect on IBI durations (p < 0.0001 and p = 0.23 for bursts and IBI, respectively; Fig. 6C).

As we found that synaptic noise by itself influences bursting, we tested the combined effect of a shorter AHP and an increased noise on burst pattern (Fig. 7). We found that this combination better mimicked the IBI duration compared to only reducing the AHP (model WT condition: 14.08 ± 0.37 for AHP + noise vs 25.39 ± 0.54 for only AHP, n = 477 and 257 bursts, respectively, p < 0.0001 and Cx-deficient: 14.66 ± 0.31, n = 250 bursts from 6 cells, p = 0.1816; Fig. 7A-B, right). In addition, it further shifted burst duration values towards small ones (model WT condition: 1.45 ± 0.013 for AHP + noise vs 1.59 ± 0.017 for only AHP, n = 477 and 257 bursts, respectively, p < 0.0001, and Cx-deficient: 1.55 ± 0.02, n = 250 bursts from 6 cells, p < 0.0001; Fig. 7A-B, left).

**Fig. 7.**
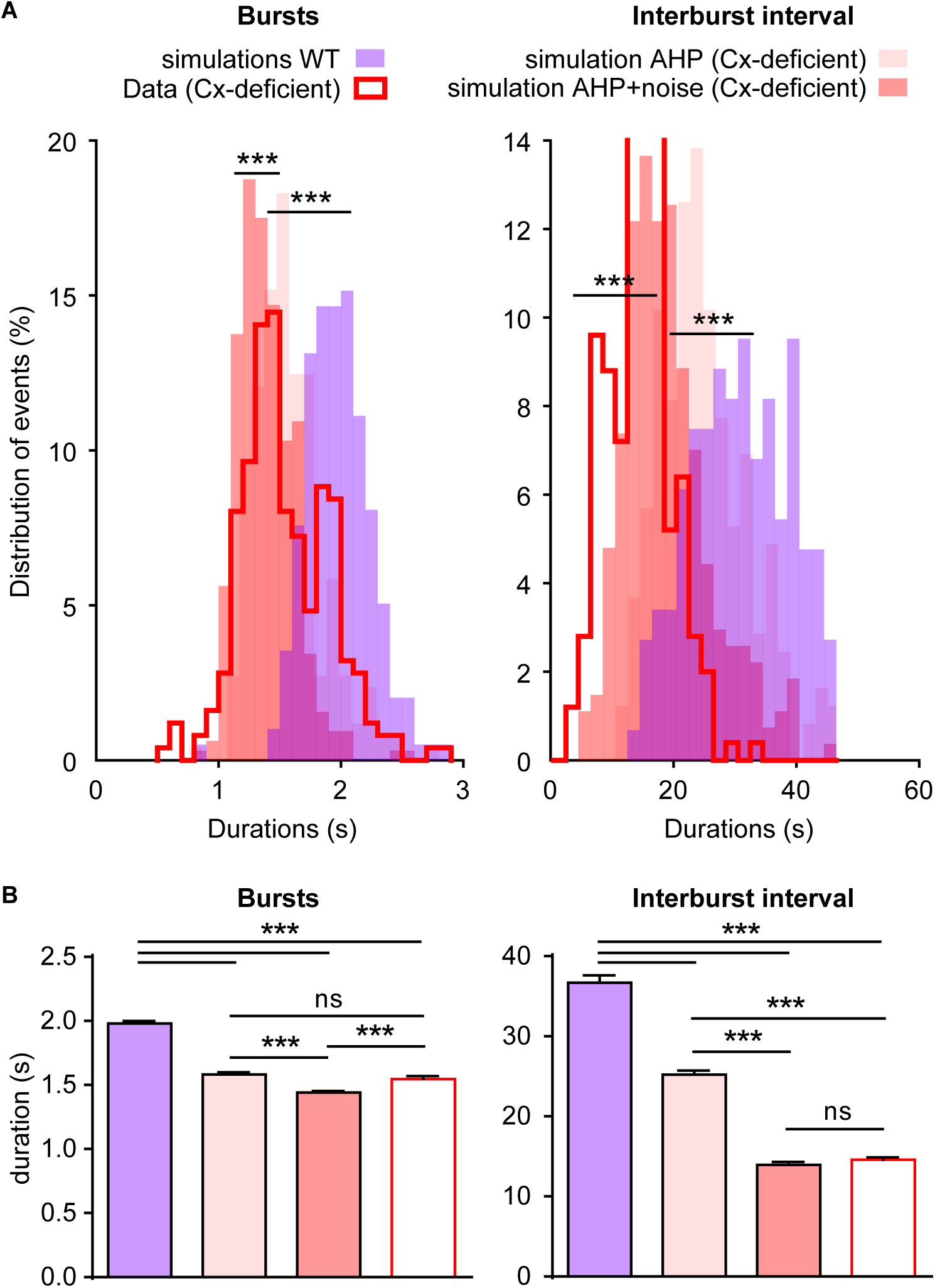
Effect of combined AHP and noise on bursting dynamics. **A**) Distribution of burst (left) and IBI (right) durations for τsAHP = 10.5 s, τmAHP = 0.35 s, TAHP = - 30 and σ = 6 (WT, light purple), for τsAHP = 5 s, τmAHP = 0.15s, TAHP = - 23 and σ = 6 (AHP levels of Cx-deficient, light rose) and for τsAHP = 5 s, τmAHP = 0.15s, TAHP = - 23 and σ = 9 (AHP and noise levels of Cx-deficient, light red), compared to the Cx-deficient data distributions (red curves) (2-sample Kolmogorov-Smirnov). T = 0 for all cases (5000 s simulations). **B**) Quantification of bursts (left) and IBI (right) durations for the conditions of panel A (ordinary one-way ANOVA with Holm-Sidak’s multiple comparisons test). Asterisks indicate statistical significance (***, p < 0.0001).

Synaptic depression is an output of the model, and depends on AHP, membrane depolarization and synaptic noise (Supplementary Fig. 4A-C). Thus, to investigate the effect of depression on burst and IBI durations, we reproduced the burst initiation depression level we observed in the WT case (Supplementary Fig. 4D, purple). To this end, we ran numerical simulations with short AHP (Cx-deficient parameters, Table1) combined with a reduced depression recovery timescale (from τ_r_ = 2.9 s to 1.9 s; Supplementary Fig. 5D, yellow). With short AHP and low depression (yellow, Supplementary Fig. 4E), we found that the bursts were longer relative to those in the Cx-deficient condition (red, short AHP and high depression; p < 0.0001), but still much shorter than in the WT case (purple, long AHP and low depression) (Supplementary Fig. 4E). This result indicates that the depression level impacts on the bursting pattern, but its change in the Cx-deficient condition does not fully account for the bursting alteration. Furthermore, AHP durations in the presence of low depression are even shorter compared to the Cx-deficient condition (p < 0.0001; Supplementary Fig. 4F), since a low depression in a neuronal network triggers more bursts (Cohen and Segal, 2011; Staley et al., 1998). These results indicate that decreasing depression in the Cx-deficient condition is not sufficient in and of itself to recover the WT phenotype and may instead be considered as another consequence of the reduced in AHP in the Cx-deficient mice. Altogether, these results indicate that the reduction in AHP is the primary factor contributing to the change in bursting pattern found in Cx-deficient mice, and that combined with an increase in synaptic noise further reproduces the burst alteration.

### Experimental validation of the role of AHP in switching between bursting patterns of wild type and Cx-deficient mice

Finally, we tested experimentally whether changes in AHP are solely responsible for the altered bursting pattern in Cx-deficient slices, as predicted by the model. To this end, we first tested in WT mice the impact of inhibiting AHP using XE-991, a blocker of voltage-gated KCNQ channels, which mediate the AHP M-current. We found that XE-991 (10 µM) significantly increased burst frequency (control 1.78 ± 0.31 vs XE-991 2.98 ± 0.420 bursts/min, p = 0.0399, n = 5 slices from 3 mice), while it reduced burst duration (control 1.68 ± 0.27 s vs XE-991 1.12 ± 0.13 s; p = 0.0340, n = 5 slices from 3 mice; Fig. 8A-B), thus mimicking alterations induced by astrocyte network disruption in Cx-deficient mice. Conversely, we found that increasing AHP in Cx-deficient slices with the anti-epileptic drug retigabine (40 µM), which activates KCNQ-type K^+^ channels, decreased burst frequency (Control, 4.75 ± 0.33 vs retigabine, 1.15 ± 0.33 bursts/min; p < 0.0001, n = 7 slices from 3 mice) and increased burst duration (Control, 1.55 ± 0.1 s vs retigabine-: 1.98 ± 0.17 s, p = 0.0481, n = 7 slices from 3 mice; Fig. 8C-D), thus rescuing WT bursting pattern in Cx-deficient mice (p = 0.2053 and 0.4672 for burst frequency and duration, respectively). In all, these results confirm the model prediction that AHP reduction primarily accounts for the altered bursting pattern observed with astrocyte disconnection. Further, they reveal that astrocyte GJ regulation of [K^+^]_e_ during bursting controls KCNQ voltage-gated K^+^ channel currents.

**Fig. 8.**
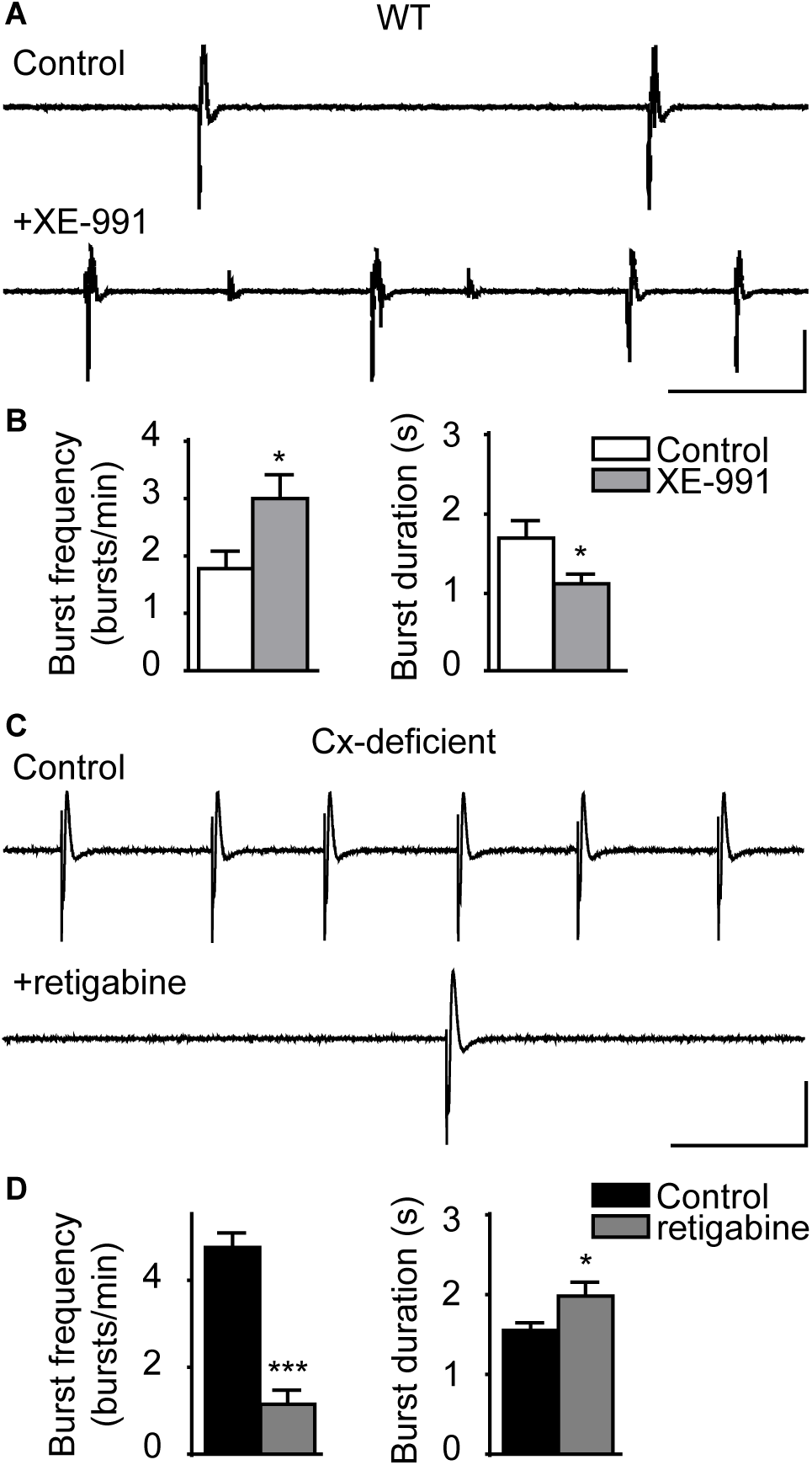
Alterations in AHP switch bursting patterns between wild type and astroglial Cx- deficient mice. **A**) Representative traces of hippocampal bursts in WT mice in control condition (top) and in the presence of XE-991 (10 µM), an inhibitor of KCNQ channels. Scale bars: 10 s, 400 μV. **B**) Quantification of burst frequency and duration (n = 5 slices from 3 mice; paired t-test). **C**) Representative traces of hippocampal bursts in Cx-deficient mice in control condition (top) and in presence of retigabine (40 µM), an activator of KCNQ channels. Scale bars: 10 s, 200 μV. **D**) Quantification of burst frequency and duration (n = 7 slices from 3 mice; paired t-test). Asterisks indicate statistical significance (*, p < 0.05; ***, p < 0.0001).

## Discussion

Here, we combined molecular, physiological and modeling approaches to investigate how astroglial networks regulate hippocampal bursting pattern. We found that astroglial GJ strengthen hippocampal population bursts primarily through [K^+^]_e_ -dependent regulation of KCNQ channels mediating AHP. Accordingly, astroglial Cx deficiency results in higher basal K^+^ levels, smaller K^+^ transients during bursting activity, and a reduced AHP, which, when restored pharmacologically by activating KCNQ-type voltage-gated K^+^ channels, rescued normal bursting patterns. These pivotal data, for the first time, identify in the central nervous system a neuronal ion channel that is directly instructed by astroglial networks to regulate network activity.

### Extracellular glutamate homeostasis, metabolic support and astroglial networks regulation of bursting pattern

In Cx-deficient mice under basal conditions, decreased glutamate clearance leads to prolonged neuronal excitatory activity via enhanced AMPAR and NMDAR activation and glutamate spillover (Pannasch et al., 2011). Here, we asked whether an impaired glutamate clearance was involved in the altered bursting pattern found in Cx-deficient slices during sustained population activity. When we partly inhibited glutamate AMPAR activation with a low dose of NBQX in Cx-deficient slices, the burst frequency was only marginally decreased, and the burst duration unchanged. In addition, when we prolonged AMPAR activation by inhibiting their desensitization with cyclothiazide in WT slices, neither the burst frequency nor duration was affected. Instead, this treatment induced spontaneous seizure-like activity, which we did not observe in Cx-deficient mice (Fig. 3). These results indicate that glutamate accumulation at the synapse is not causing the bursting phenotype of Cx-deficient mice.

In the absence of functional astroglial networks, decreased presynaptic release probability was observed during spontaneous bursting activity (Chever et al., 2016). Remarkably, a decreased release probability was also reported during active states of physiological slow oscillations (Crochet et al., 2005). Consistently, action potential-induced synchronous synaptic release of glutamate initiate bursts that are sustained until vesicle pools are exhausted (Cohen and Segal, 2011; Staley et al., 1998). Accordingly, we here found in Cx-deficient mice that the readily releasable pool of synaptic vesicles during bursting was decreased, as shown by the impaired responses to prolonged 10 Hz stimulation (Fig. 2E-G). As previously reported, the enhanced synaptic bombardment that causes neuronal depolarization and spontaneous firing between bursts likely decreases the readily releasable pool of synaptic vesicles during bursting, thus reducing synaptic efficacy, neuronal synchronization and burst strength (Chever et al., 2016).

GJ-mediated astroglial networks sustain glutamatergic activity by providing lactate to active neurons in an activity-dependent manner (Rouach et al., 2008). Impaired energy supply to neurons in Cx-deficient mice could therefore contribute to the reduced burst duration and neuronal synchronization. However, our recordings were performed in 11 mM glucose-containing ACSF, thereby bypassing a possible reduction in glucose supply as an explanation for the Cx-deficient burst phenotype. In addition, we found in WT mice that extracellular glucose deprivation halved the burst frequency after 20 min and almost fully blocked activity after 30 min, an effect which did not reproduce the Cx-deficient bursting pattern (Fig. 4). Hence the altered bursting pattern in astroglial Cx deficient mice is unlikely to result from an impaired energy supply.

### Gap junction-mediated astroglial networks modulate extracellular potassium during bursting

[K^+^]_e_, tightly regulated in the brain and kept close to 3 mM in basal conditions, undergoes local changes upon neuronal activity. Given the small volume of the extracellular space and the low baseline [K^+^]_e_, limited neuronal K^+^ efflux may actually induce significant changes in [K^+^]_e_. Physiological neuronal activity leads to [K^+^]_e_ increases of less than 1 mM, while pathological activity, such as seizures, can increase [K^+^]_e_ up to 10-12 mM (Bellot-Saez et al., 2017). Changes in [K^+^]_e_ can strongly impact several neuronal processes, including voltage-gated ion channel activity, synaptic transmission, neurotransmitter transport, and membrane potential maintenance and excitability. Rapid changes in [K^+^]_e_ are thus tightly controlled by passive diffusion and cellular K^+^ clearance mechanisms (Walz, 2000). Effective removal of [K^+^]_e_ is indeed vital for maintaining brain homeostasis and limiting neuronal network hyperexcitability during physiological brain processes. Early on, GJ-connected retinal glial cells were recognized to play a role in controlling [K^+^]_e_ through spatial buffering (Karwoski et al., 1989). Local excess of K^+^, dispersed through interconnected glial cells, can transfer K^+^ ions from sites of elevated [K^+^]_e_ over relatively long distances to sites with lower [K^+^]_e_.

Astroglial GJ have been assumed to play a similar role. However, studies in astroglial Cx-deficient mice suggest that astroglial GJ only partially account for K^+^ spatial buffering in the hippocampus. GJ-mediated currents indeed represent ∼30 % of the astrocyte whole-cell currents (Wallraff, 2006), and although they contribute to [K^+^]_e_ homeostasis at both physiological (single and paired-pulse stimulation) (Pannasch et al., 2011; Wallraff, 2006) and pathological levels (trains of stimulations) (Wallraff, 2006), Cx-deficient astrocytes still display a large K^+^ clearance capacity. More recent work found that acute pharmacological inhibition of GJs does not alter synaptically-evoked [K^+^]_e_ transients in hippocampal slices, but only increases large and localized K^+^ variations exceeding ∼10 mM (Breithausen et al., 2020). However, this study used carbenoxolone, which is not a specific blocker of astroglial GJ, as it blocks GJ from all cell types, and is a mineralocorticoid agonist with several other off-target effects, inhibiting numerous ion channels and pumps such as Na/K ATPases (Zhou et al., 1996), chloride channels (Böhmer et al., 2001), voltage-gated Ca^2+^ channels (Vessey et al., 2004), pannexin1 channels (Michalski and Kawate, 2016) as well as neurotransmitter receptors such as AMPARs (Tovar et al., 2009) and GABA_A_Rs (Ransom et al., 2017).

Carbenoxolone thus alters intrinsic neuronal membrane properties (Rouach, 2003) and synaptic transmission (Vessey et al., 2004), besides having some neurotoxic effects (Rouach, 2003). These multiple actions of carbenoxolone, independent of GJ inhibition, thus preclude its use to assess specific impacts of astroglial GJ on [K^+^]_e_.

Given the current lack of a specific pharmacological inhibitor of astroglial GJ communication and the pleiotropic effects of [K^+^]_e_, we here combined molecular and modeling approaches to study the impact of astroglial GJ on [K^+^]_e_ and network activity. Based on previous work showing that an increment of [K^+^]_e_ from 5 to 10 mM can cause a 5- fold increase in the frequency of hippocampal interictal events (Rutecki et al., 1985; Traynelis and Dingledine, 1988), we hypothesized that the increased bursting activity in mice with disconnected astrocytes results from K^+^ accumulation in the extracellular space. Further, since disconnected astrocytes display increased volume associated with decreased extracellular space (Pannasch et al., 2011), K^+^ clearance by extracellular diffusion is also likely to be impaired, and thus contribute to extracellular K^+^ build-up. Accordingly, we here found that resting [K^+^]_e_ between bursts is increased compared to WT mice (Fig. 1A-B) (Chever et al., 2016). This is in agreement with the steady-state depolarization of CA1 pyramidal cells (Chever et al., 2016) and astrocyte membrane potentials (Fig. 1D-E) in a regime of bursting activity in mice with disconnected astrocytes. It is noteworthy that this effect is specific for the bursting regime of activity, as it was not observed in basal conditions (Pannasch et al., 2011). In agreement with these observations, we found that K^+^ transients during bursts are smaller and shorter in mice with disconnected astrocytes compared to WT mice, as well as burst-associated or synaptically-evoked astrocytic membrane depolarizations, which represent a sensitive measure of [K^+^]_e_ levels during activity (Fig. 1).

Astrocytic disconnection indeed prevents optimal coordination of neuronal populations and impairs synchronization (Chever et al., 2016). Astrocytes, as part of the tripartite synapse, modulate neuronal synchronization and network activity, by preventing excessive accumulation of K^+^ (Bellot-Saez et al., 2017). Furthermore, alterations in [K^+^]_e_ due to astrocyte disconnection may also alter local electric fields and synaptic current waveforms, thereby impacting on the whole network signal integration (Amiri et al., 2013) and mediating transitions between tonic and phasic neuronal oscillations (Frohlich et al., 2006). Remarkably, we also found that changing resting K^+^ levels induced WT-like bursting patterns in astroglial Cx-deficient mice and vice versa, which points to the contribution of impaired [K^+^]_e_ homeostasis in the altered network activity of mice with disconnected astrocytes. Thus, the control of [K^+^]_e_ by GJ-connected astrocytes not only modulate neuronal excitability at the cellular level, but also regulate the global activity of neuronal networks.

### KCNQ channels are a target of astroglial gap junctions

Given that K^+^ acts on multiple membrane (membrane potential and AHP) and synaptic properties (noise and depression), that are all altered in mice with disconnected astrocytes ((Chever et al., 2016) and Fig. 2), deciphering the physiological target of [K^+^]_e_ involved in the astroglial control of bursts using experimental approaches is challenging. Hence, we here developed a novel neuronal network model based on experimental data and underlying burst generation. In this model, which accounts for synaptic facilitation/depression (Dao Duc et al., 2015; Holcman and Tsodyks, 2006; Markram et al., 1998), we integrated experimental data on membrane properties and [K^+^]_e_ dynamics, and introduced an AHP component, without making a distinction between specific K^+^ channels. To calibrate the model, we identified parameters by minimizing the difference between the distribution of bursts and IBI durations from simulations and experiments. This approach predicted that the astroglial network regulation of [K^+^]_e_ sets bursting pattern by controlling AHP. Interestingly, the changes of AHP parameters (τ*_sAHP_*, τ*_mAHP_* and T*_AHP_*) needed in the model to reproduce the burst and interburst duration alteration observed with Cx deficiency are similar to those reported in our previous work (Chever et al., 2016). Indeed, experimentally we observed that neuronal AHP duration is halved and its amplitude is reduced by ∼ 40% when astrocytes are disconnected (Chever et al., 2016). Similarly, in our model we reproduced Cx-deficiency-dependent burst alteration with a 50 % decrease of τ*_sAHP_* and τ*_mAHP_*, which relates to AHP duration, and a ∼ 30 % reduction of T*_AHP_*, which reflects AHP amplitude.

This suggested that impaired membrane depolarization and repolarizations of neurons in mice with disconnected astrocytes altered the proper temporally and spatially-restricted functioning of voltage-gated ion channels during bursting. Our electrophysiological recordings confirmed the role of AHP in the astroglial network regulation of bursting pattern, and identified that GJ-mediated control of AHP results from modulation of KCNQ voltage- gated K^+^ channel currents.

Interestingly, loss of Cx43 GJ channels in the heart causes a decrease in action potential duration in myocytes by increasing sustained repolarizing and inward rectifier K^+^ currents (Danik et al., 2008). Furthermore, Cx43 deletion in adult ventricular and fetal atrial myocytes also decreases the amplitude of the Na_v_1.5-mediated sodium current (Desplantez et al., 2012; Jansen et al., 2012), thereby indicating that Cx43 channels are necessary for proper sodium current function. Remarkably, it has also been shown in pancreatic β-cells that Cx36 containing GJ channels coordinate K_ATP_ channel activity to promote synchronized and oscillatory insulin secretion under stimulatory levels of glucose or global β-cell inhibition at basal glucose level (Farnsworth and Benninger, 2014). Consistent with these data, we show that in the central nervous system, astroglial GJ modulate neuronal excitability primarily via [K^+^]_e_ regulation of AHP mediated by KCNQ channels. It is noteworthy that the regulation of AHP/KCNQ channels by [K^+^]_e_ has been reported in different systems. Increase in [K^+^]_e_ reduces AHP in hypoglossal motoneurons (Viana et al., 1993) and KCNQ2/3 currents in HEK cells (Prole et al., 2003). This latter inhibitory effect has been proposed to result from binding of K^+^ ions within the outer pore region of the channel (Doyle et al., 1998; Zhou et al., 2001), which causes a physical occlusion of the outer pore or disturbance of K^+^ binding sites along the pore axis (Prole et al., 2003). In addition to this K^+^-mediated inhibition, the increased [K^+^]_e_ reduces the driving force guiding K^+^ ions through the channels. Together, these effects likely result in a reduction of KCNQ2/3 currents, thus shortening AHP, increasing excitability and facilitating burst triggering in Cx-deficient mice.

By identifying KCNQ channels as downstream/molecular targets of astroglial GJ, our data uncover a previously unknown molecular mechanism underlying astroglial network regulation of bursting pattern with therapeutic potential in CNS disease.

## Methods

### Experimental design

The main objective of this study was to explore by which mechanism gap junction-mediated astrocytic networks modulate neuronal bursting patterns in mouse hippocampal slices. First, we investigated experimentally the physiological mechanism underlying astrocyte-mediated regulation of neuronal network activity. Subsequently, we used a modeling approach to predict the underlying neuronal molecular target. Finally, we validated experimentally the model prediction.

### Animals

Experiments were carried out according to the guidelines of the European Community Council Directives of January 1^st^ 2013 (2010/63/EU) and all efforts were made to minimize the number of used animals and their suffering. Experiments were performed in the hippocampus of wild-type mice (WT) and Cx30^-/-^Cx43^fl/fl^ hGFAP-Cre mice (Cx-deficient), provided by Pr. K. Willecke (University of Bonn, Germany), with conditional deletion of Cx43 in astrocytes and additional total deletion of Cx30 (Wallraff, 2006). For all analyses, mice of both genders and littermates were used and *ex vivo* slice electrophysiology was performed on P16-P25 mice as previously described (Rouach et al., 2008).

### In vitro slice electrophysiology

Acute transverse hippocampal slices (400 μm) were prepared as previously described (Rouach et al., 2008) from 16 to 25 days old WT and Cx-deficient mice. Slices were maintained in a storage chamber containing a standard artificial cerebrospinal fluid (ACSF; 119 mM NaCl, 2.5 mM KCl, 2.5 mM CaCl_2_, 1.3 mM MgSO_4_, 1 mM NaH_2_PO_4_, 26.2 mM NaHCO_3_, and 11 mM glucose, saturated with 95% O_2_ and 5% CO_2_) for 30 minutes, then stored in a magnesium-free ACSF in the presence of picrotoxin (100 μM) for at least 1 h before recording so that the slices can spontaneously generate population bursts. Slices were transferred in a submerged recording chamber mounted on an Olympus BX51WI microscope equipped for infrared-differential interference (IR-DIC) microscopy and were perfused with 0 Mg^2+^-picrotoxin ACSF at a rate of 2 ml/min. Extracellular field and whole-cell patch-clamp recordings of astrocytes were performed. Field excitatory bursts were recorded with glass pipettes (2–5 MΩ) filled with ACSF and placed in *stratum radiatum*. Prolonged repetitive stimulation was performed at 10 Hz for 30 s. Whole-cell recordings were obtained from visually identified CA1 *stratum radiatum* astrocytes using 5-10 MΩ glass pipettes filled with 105 mM K-gluconate, 30 mM KCl, 10 mM HEPES, 10 mM phosphocreatine, 4 mM ATP-Mg, 0.3 mM GTP-Tris, and 0.3 mM EGTA (pH 7.4, 280 mOsm). The stimulations consisted of 0.1 and 0.5 ms electrical pulses (15 μA) applied through a glass pipette located in the *stratum radiatum*. CA1 astrocytes resting membrane potential, membrane and series resistance as well as membrane capacitance were monitored throughout the recordings.

Measurements of membrane resistance and capacitance were performed on astrocytes clamped at - 80 mV. Responses (neuronal fEPSP slope) to repetitive stimulation (10 Hz, 30 s) were binned (bin size 1.2 s) and normalized to mean baseline responses measured at 0.1 Hz before repetitive stimulation. Field potentials and patch-clamp recordings were acquired with Axopatch-1D amplifiers (Molecular Devices, USA), digitized at 10 kHz, filtered at 2 kHz, stored and analyzed on computer using pCLAMP9 and Clampfit10 software (Molecular Devices, USA).

For multi-electrode array (MEA) recordings, hippocampal slices were transferred on planar MEA petri dishes (200-30 ITO electrodes, organized in an 12x12 matrix, with internal reference, 30 µm diameter and 200 µm inter-electrode distance; Multichannel Systems, Germany). They were kept in place by using a small platinum anchor. The slices on MEAs were continuously perfused at a rate of 2 ml/min with a magnesium-free ACSF containing picrotoxin (100 µM), as previously described (Chever et al., 2016). Pictures of hippocampal slices on MEAs were used to identify the location of the electrodes through the different hippocampal regions and to select the electrodes of interest. Data were sampled at 10 kHz and network spontaneous activity was recorded at room temperature by a MEA2100-60 system (bandwidth 1-3000 Hz, gain 2x, Multichannel Systems, Germany) through the MC Rack 4.5.1 software (Multichannel Systems, Germany).

### Preparation of K^+^-sensitive microelectrodes and measurement of extracellular K^+^ concentration

Single-barreled K^+^-selective microelectrodes were prepared using thin-walled borosilicate capillaries (GC150T-7.5, Harvard Apparatus, USA). The interior walls of the capillaries were silanized with silan vapors (*N*,*N*-Dimethyltrimethylsilylamine, Sigma Aldrich, France) for 15 min and dried at 200°C for 100 min. The tip of K^+^-selective microelectrodes was filled with Potassium ionophore I (Cocktail A, Sigma Aldrich, France) and the rest of the electrode was backfilled with 200 mM KCl in ACSF background. The reference electrode was made of standard patch-clamp glass (GC150F-10, Harvard Apparatus, USA) and filled with ACSF containing (in mM): 119 NaCl, 2.5 KCl, 1 NaH_2_PO_4_, 26.2 NaHCO_3_, 2.5 CaCl_2_, 1.3 MgSO_4_ and 11 glucose. The K^+^-selective microelectrodes were calibrated using solutions containing 0.5, 1, 2, 3, 4, 5, 6, 8, 10, 12, 20 and 40 mM KCl. The extracellular K^+^ concentration was recorded in WT and Cx-deficient mouse hippocampal slices in a recording chamber mounted on an Olympus BX51WI microscope equipped for infrared differential interference microscopy and with 40x objective; slices were perfused with standard ACSF or magnesium-free ACSF containing picrotoxin (100 µM) at a rate of 2 ml/min. Both the K^+^-selective and reference electrodes were placed in CA1 *stratum radiatum*, so that their tips were within 10 μm distance from each other. Data were acquired using Axopatch 200B amplifier, sampled at 20 kHz, low pass filtered (2 kHz), digitized (Digidata 1440), and stored and analyzed on computer using pCLAMP 9 and Clampfit 10 software (all from Molecular Devices, USA). The signal from the reference electrode was offline subtracted from the signal of the K^+^- selective electrode to obtain a signal proportional to actual K^+^ concentration. The relationship between the measured voltage and the actual K^+^ concentration was derived from the log-linear fit function.

### Burst Analysis

Raw data were analyzed with MC Rack (Multi-Channel System, Reutlingen, Germany). Detection of bursts was performed using the “Spike Sorter” algorithm, which sets a threshold based on multiples of standard deviation of the noise (5-fold) calculated over the first 500 ms of recording free of electrical activity. A 5-fold standard deviation threshold was used to automatically detect each event, which could be modified in real-time by the operator on visual check if needed. Analysis of burst duration was performed using Neuroexplorer (version 4.109, Nex Technologies, USA).

### Drugs

NBQX and cyclothiazide were from Tocris (UK); all the other products were obtained from Sigma-Aldrich (France).

### Generalized depression-facilitation model accounting for potassium dynamics

The facilitation-depression model (Dao Duc et al., 2015; Tsodyks and Markram, 1997) is a mean-field type representation of a sufficiently connected neuronal network. It consists of three equations for the mean voltage *h* (or firing rate), the mean depression *y*, and the mean synaptic facilitation *x*. However, this neuronal-network model does not account for long hyperpolarization periods, due to K^+^ channel activation (McKiernan and Marrone, 2017), leading to a refractory period. To account for these periods, we modified this initial model by introducing two new features:

1) A hyperpolarized equilibrium state, in addition to the resting state. This new state is defined by a negative value for the voltage h, with a new threshold (*T*_AHP_ = -30, see Table 1) in addition to *T*_0_ = 0
2) A medium and slow recovery of the voltage during AHP to the resting potential, modeled by two time constants τ*_mAHP_* and τ*_sAHP_*.

The general system becomes

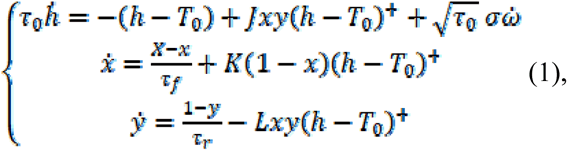

where, 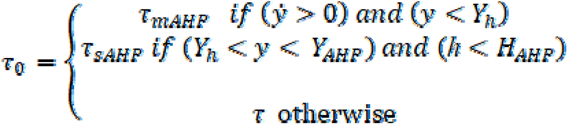 and 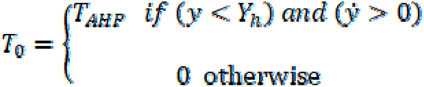

The threshold linear function *h*^+^ = max(*h*,0) represents the firing rate and is a Gaussian white noise centered at 0 and of variance 1. The noise amplitude is σ. The parameters J, K, L represent the synaptic connectivity, the facilitation increases during bursts and the rate of the vesicular release probability (i.e. the depression) decays (Bart et al., 2005; Holcman and Tsodyks, 2006). The time constants τ_f_ (facilitation) and τ_r_ (depression recovery) are defined in Table 1.

The equations of the model recapitulate the burst decomposition presented in Fig. 5A. Specifically, we define four different phases:

1) The burst initiation is triggered by a voltage fluctuation, modeled by a Gaussian white noise of low amplitude σ, when the dynamical system (eq. 1) is at equilibrium (T_0_ = 0 and τ_0_ τ). This phase is dominated by fast spiking dynamics (τ_0_ τ) that occurs during neuronal bursting (Fig. 5A, Step 1, blue).
2) The bursting phase ends when the depression variable y reaches its minimum (i.e. when the readily releasable pool is depleted). This timepoint marks the beginning of the hyperpolarization phase with a new time scale τ_0_ τ_mAHP_ and a new resting value T_0_ T_AHP_ < 0 for the variable h. In practice, this phase is defined by the condition y < Y_h_ and (y’ > 0), Step 2 (red in Fig. 5A). These changes force the voltage to hyperpolarize.
3) The third phase defines the slow recovery to equilibrium: it is characterized by a new time constant τ_0_ τ_sAHP_, the slowest time scale, and the resting value is now T_0_ 0 (Step 3, purple in Fig. 5A). When the conditions y > Y_AHP_ and h ≥ h_AHP_ are satisfied, the dynamics is governed again by the fast timescale τ_0_ τ and a resting state T_0_ T 0 (Fig. 5A, Step 4). This period lasts until y > Y_AHP_ or h > h_AHP_ (i.e. when the system is back at its equilibrium).
4) The last phase (QP, green) is identical to the first phase and represents the periods of small fluctuations around the equilibrium of the system until the next burst is generated. The role of the noise is to generate spontaneous bursts (Fig. 5B). The parameters K, L and J are adapted from to fit the MEA data (see Table 1). The fine-tuning of all parameters is obtained by best fitting the burst and IBI time distribution, as described in the Results section. The AHP parameters are determined using the order of magnitudes observed in CA1 hippocampal pyramidal neurons (McKiernan and Marrone, 2017).

### Numerical Simulations

We ran numerical simulations of equations (1) in MATLAB using a Runge-Kutta 4 scheme that we implemented, with *δt* = 10 ms (we also used *δt* = 1 *ms* to confirm the robustness of the numerical scheme and we could obtain similar results using Euler’s scheme with a small *δt*).

### Time series segmentation

To detect the bursts and IBI, we developed a segmentation procedure based on various time series: MEA recordings of hippocampal slices from WT and Cx-deficient mice, patch-clamp recordings of hippocampal pyramidal cells from WT and Cx-deficient mice and numerical simulations. The method is detailed in the Supplementary Information, but we here briefly summarize the principles used for the experimental data: we first filtered the individual action potentials using a sliding window of length *T_w_* =1*s* to compute the local average 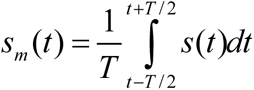. A burst is detected when *s_m_* (*t*) exceeds a threshold and the end of the burst is determined when *s_m_* (*t*) reaches its equilibrium value . This segmentation is used to extract the burst and IBI durations in both MEA and patch-clamp recordings (see supplementary information for details). The threshold is estimated by fitting a horizontal line to the epochs preceding the burst, and we define the burst detection threshold as 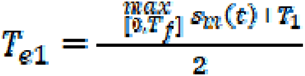.

### Statistical analysis

Data are expressed as mean ± SEM, unless otherwise stated. Statistical significance for between groups comparisons was determined by paired and unpaired two tailed t-tests with Welsh’s correction when populations had different standard deviations or non-parametric Mann-Whitney tests when data were not normally distributed. One-way ANOVA with Dunnett *post hoc* test was performed for 0 glucose experiment. Repeated measures two-way ANOVA was performed for repetitive stimulation (10 Hz, 30 s). Two-sample Kolmogorov-Smirnov test was used for distribution comparison. Differences were considered significant at p < 0.05. Statistical analysis was performed using GraphPad Prism 5 software and figures were prepared using Adobe Illustrator CS3. Exact p values are given unless p < 0.0001 or p > 0.9999.

## Acknowledgements Funding

European Research Council (Consolidator grant #683154) (NR)

European Union’s Horizon 2020 research and innovation program (Marie Sklodowska- Curie Innovative Training Networks, grant #722053, EU-GliaPhD) (NR)

Ligue Francaise contre l’Epilepsie (ED) Aviesan (ED)

Grant Agency of the Czech Republic (GACR: 19-02046S) (LV, HP) French government’s scholarship program (HP)

Fondation pour la Recherche Medicale (OC)

French Research Ministry (ED386, Ecole Doctorale de Sciences Mathématiques Paris centre) (LZ)

Fondation pour la Recherche Medicale (FRM FDT202012010690) (LZ)

## Author contributions

Conceptualization: ED, DH, NR

Patch-clamp experiments and analysis: ED, LZ, OC

MEA experiments and analysis: ED

K^+^ sensitive microelectrodes: LV, HP

Mathematical modeling: LZ, DH

Data interpretation: ED, NR

Writing: ED, LZ, DH, NR

## Data and materials availability

All data are available in the main text or the supplementary materials and they are available from the corresponding author upon reasonable request. Material requests and correspondence should be addressed to N.R. and D.H.

## Supplementary Materials

Supplementary Information is available for this paper:

Supplementary Methods Figs. S1 to S4

## Supplementary Methods

### Signal processing

#### Noise analysis

To quantify the level of the noise in the membrane potential of hippocampal pyramidal cells from wild type (WT) and astroglial Cx-deficient mice, we selected the time periods outside bursting and AHP (quiescent phase) in the patch-clamp recordings. To obtain a significant time period, we concatenated these phases for different cells that were rarely spiking.

We computed the power spectrum for the WT and Cx-deficient voltage time series (1000 s for the WT and 600 s for the Cx-deficient), and filtered them using a low pass filter (Butterworth of order 3) with a cutoff frequency of 100 Hz that we denote *P*_WT_ and *P*_Cx-deficient_. To extract from these power spectrums further information, we used the model (1, main text) around its equilibrium point in the presence of a continuous Brownian noise. Under the condition of neither burst nor AHP, the voltage equation can be approximated by:

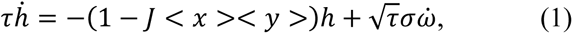

where we use the approximation that the facilitation and depression are constant equal to <x> and <y> respectively. Equation (1) is an Ornstein-Uhlenbeck process and the associated power spectrum is a Lorentzian:

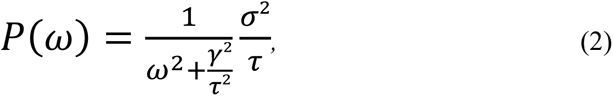

where *γ* = 1−*J*<x><y>. We fitted equation (2) to the filtered power spectrums *P*_WT_ and *P*_Cx-deficient ,_ as shown in Fig. S1C. The amplitude of the noise for the wild type case is 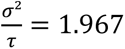 and for the Cx-deficient case 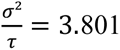. We conclude that the amplitude of the noise is doubled in Cx-deficient compared to WT mice. Interestingly, the cutoff frequency *γ* is similar 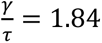 in WT and 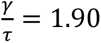 in Cx-deficient mice.

#### Time-series segmentation

We describe here the segmentation methods that we developed to differentiate bursting and AHP from resting phases in the case of patch-clamp and MEA recordings and numerical simulations.

• For the patch-clamp data, we first apply a low-pass filter to the input membrane potential *s*(*t*) using a sliding time window of length *T_w_* = 1 *s* resulting in the output signal *s_m_*(*t*) (Fig. S4 A1). We detect burst initiation when the filtered signal reaches a threshold such as *s_m_*(*τ^i^*) = *T_e_*_1_, where 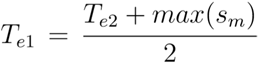 mV is the average between the maximum value of the filtered signal and the resting membrane potential *T_e_*_2_ (Fig. S3 A2, yellow dotted line). We note that although burst initiation occurred before *τ^i^* when the signal *s_m_* was at resting state, we neglected this delay of the order of few ms. We determine the time of burst termination when the signal decreases to its resting state defined by *s_m_*(*τ^e^*) = *T_e_*_2_ (Fig. S3 A2, red line). The end of the burst *τ^e^* is the beginning of the hyperpolarization phase that lasts until the signal *s_m_* has increased back to its resting value *T_e_*_2_ at time *τ^a^* (Fig. S3 A3, magenta trace). Finally, we define a quiescent phase as the time period between the end of hyperpolarization until the initiation of the next burst (Fig. S3 A3, green trace). In summary, we segmented the *n^th^* burst duration 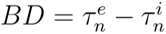 (Fig. S3, blue line), hyperpolarization duration 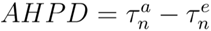, (magenta line), and quiescent phases 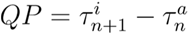 (green line).
• For MEA recordings, we apply a low-pass filter to the input field recordings using a sliding time window of length T_w_ = 0.4 s. We detect bursts and IBI on the absolute value of the filtered signal |s_m,MEA_|. Specifically, we detect the burst initiation time τ_i_ when the signal reaches one third of its maximum value such as 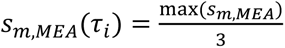 and the end time τ_e_ of the burst when 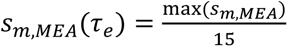 Because it is not possible to visualize the AHP in MEA recordings, this segmentation only provides the statistics of burst and IBI durations.
• In numerical simulations, the burst initiation is detected when the mean voltage *h*(*τ*) = *T*_1_ (Fig. S3 B1). Here we cannot neglect the time delay between the burst initiation and the time detection *τ*. We set the time *τ^i^* of burst initiation as the last time previous to *τ* where the mean voltage *h* was equal to its resting value: *h*(*τ^i^*) = *T* (Fig. S3 B2). Similarly, we detect burst termination by finding the time *τ* such as the mean voltage passes a second threshold *T*_2_ *< T* leading to *h*(*τ*) = *T*_2_. To account for a possible delay, we consider as the burst termination the last time *τ^e^* before *τ* where *h* is equal to its resting value *h*(*τ^e^*) = *T* (Fig. S3 B2). Similarly, to the patch recordings, we detect the hyperpolarization and quiescent phases.

The distributions obtained from this segmentation are given in Fig. S1 A-B.

## Supplementary Figures

**Fig. S1.**
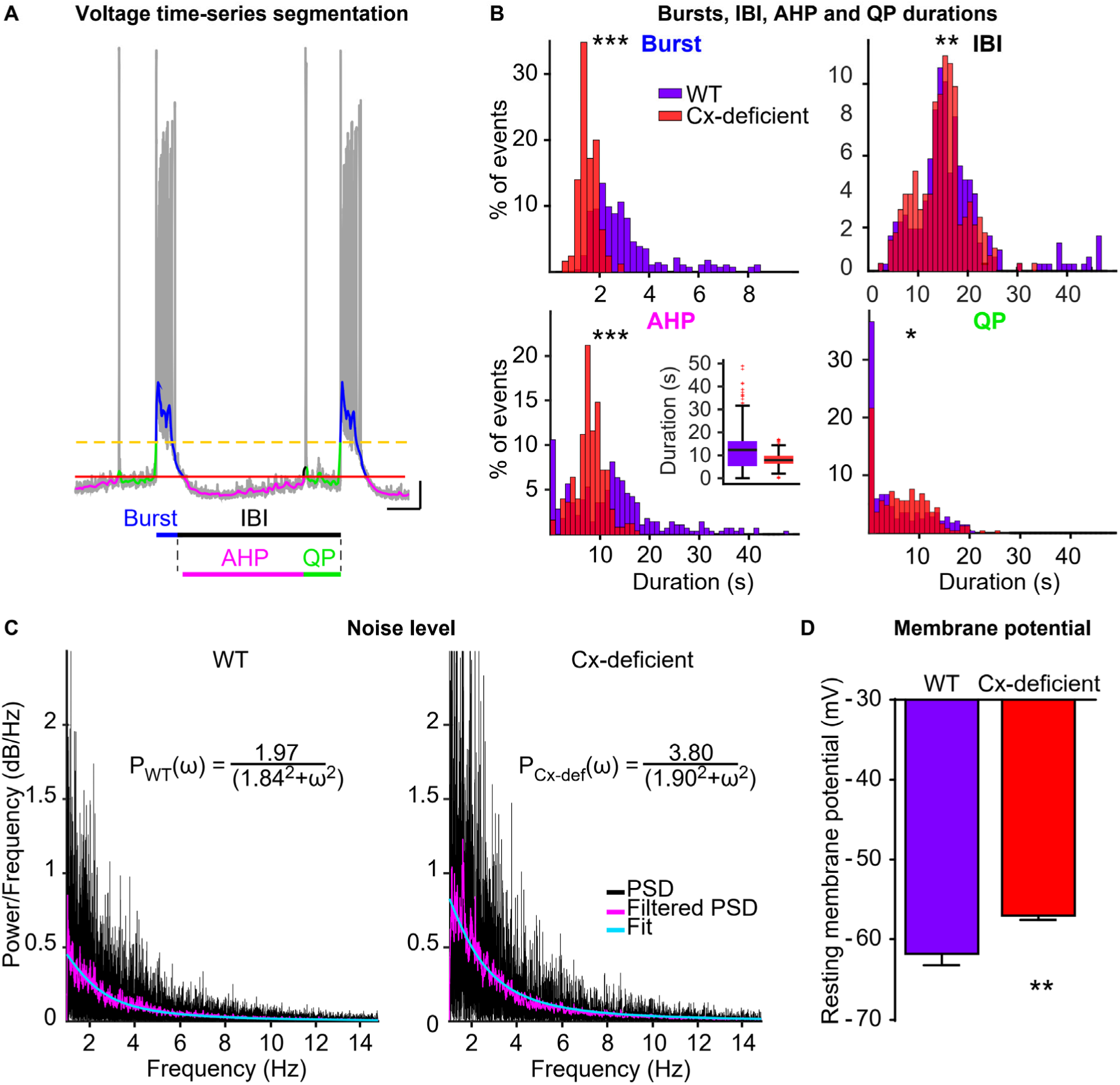
Statistics of burst dynamics from electrophysiological time-series in neurons from WT and Cx-deficient mice. **A**) Segmentation of patch-clamp recordings in hippocampal neurons in 3 phases: burst (blue), AHP (pink), and QP (green) using two thresholds T_1_ and T_2_ (green and yellow dotted lines). AHP and QP form IBI (black). Scale bar: 10 s, 10 mV. **B**) Distributions of bursts (upper left), IBI (upper right), AHP (lower left, inset boxplots) and QP (lower right) durations from neurons in WT (purple) and Cx-deficient (red) mice (n = 10 neurons for WT and n = 6 neurons for Cx-deficient; two sample t- test). **C**) Power spectrum (black) of the QPs and power spectrum P_F_ for the filtered signal at fc = 30 Hz (low-pass filter, pink trace). The fit of a Lorentzian 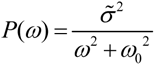 for the WT (left) and Cx-deficient (right) is shown in blue. The extracted noise amplitude *σ̃*-^2^is 1.97 and 3.80 for WT and Cx-deficient mice, respectively. **D**) Resting membrane potentials in neurons from WT (n = 12 cells, purple) and Cx-deficient mice (n = 10 cells, red; p = 0.0075, unpaired t-test). Asterisks indicate statistical significance (**, p < 0.01; ***, p < 0.0001).

**Fig. S2.**
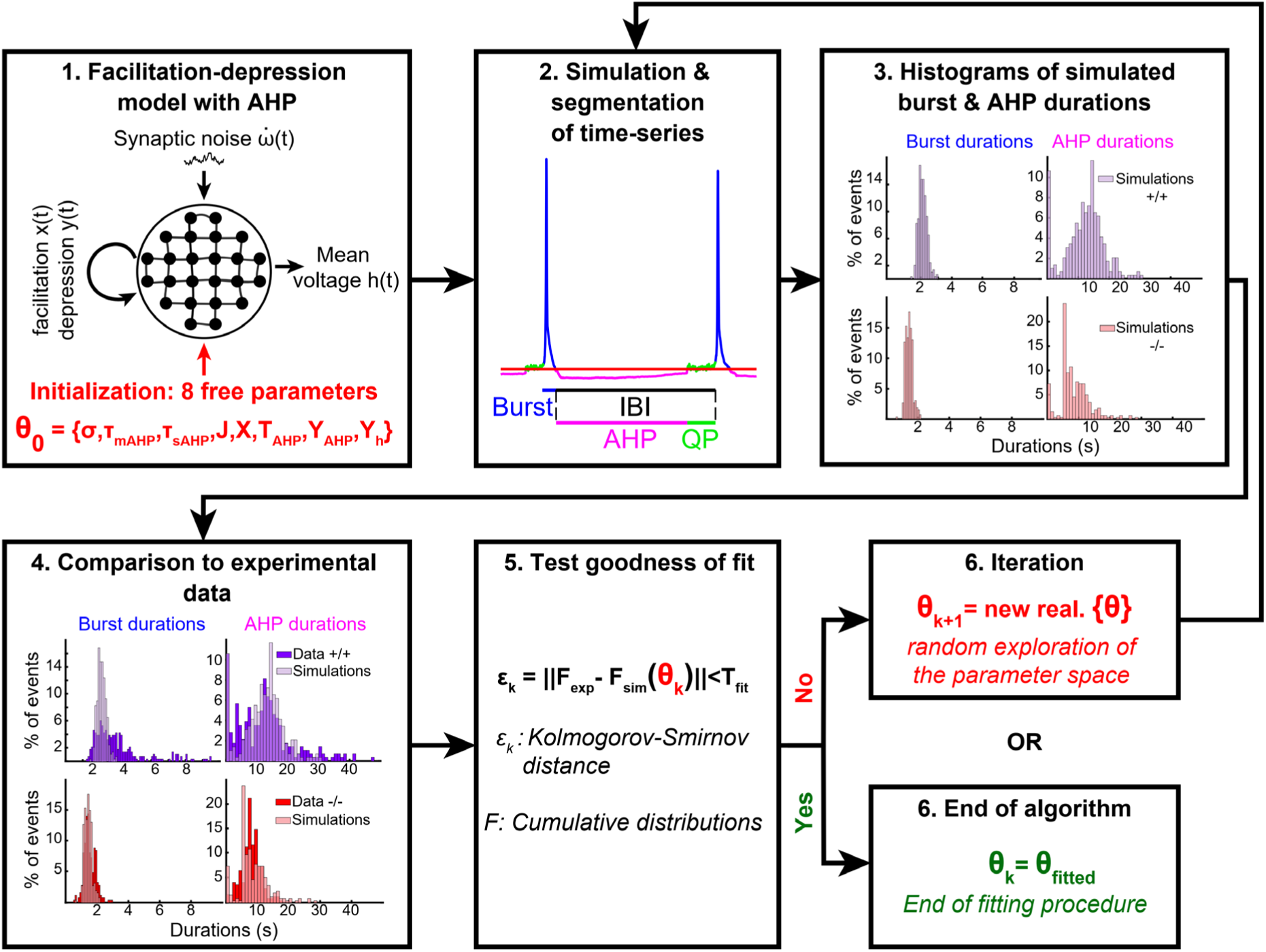
Model fitting pipeline. (1) The model has 8 free parameters (vector **θ**) that need to be fitted. (2) At each step **k**, run and segment 5000s simulation of time series with the parameters **θ_k_**. (3) Build the histograms of burst and AHP durations from the numerical simulations. (4-5) Compare the simulated histograms to the experimental ones by computing the Kolmogorov-Smirnov distance ɛ_k_ between the experimental and simulated distributions. (6) If ɛ_k_ is higher than the threshold T_fit_ then we iterate at step (2) otherwise we terminate the fitting algorithm.

**Fig. S3.**
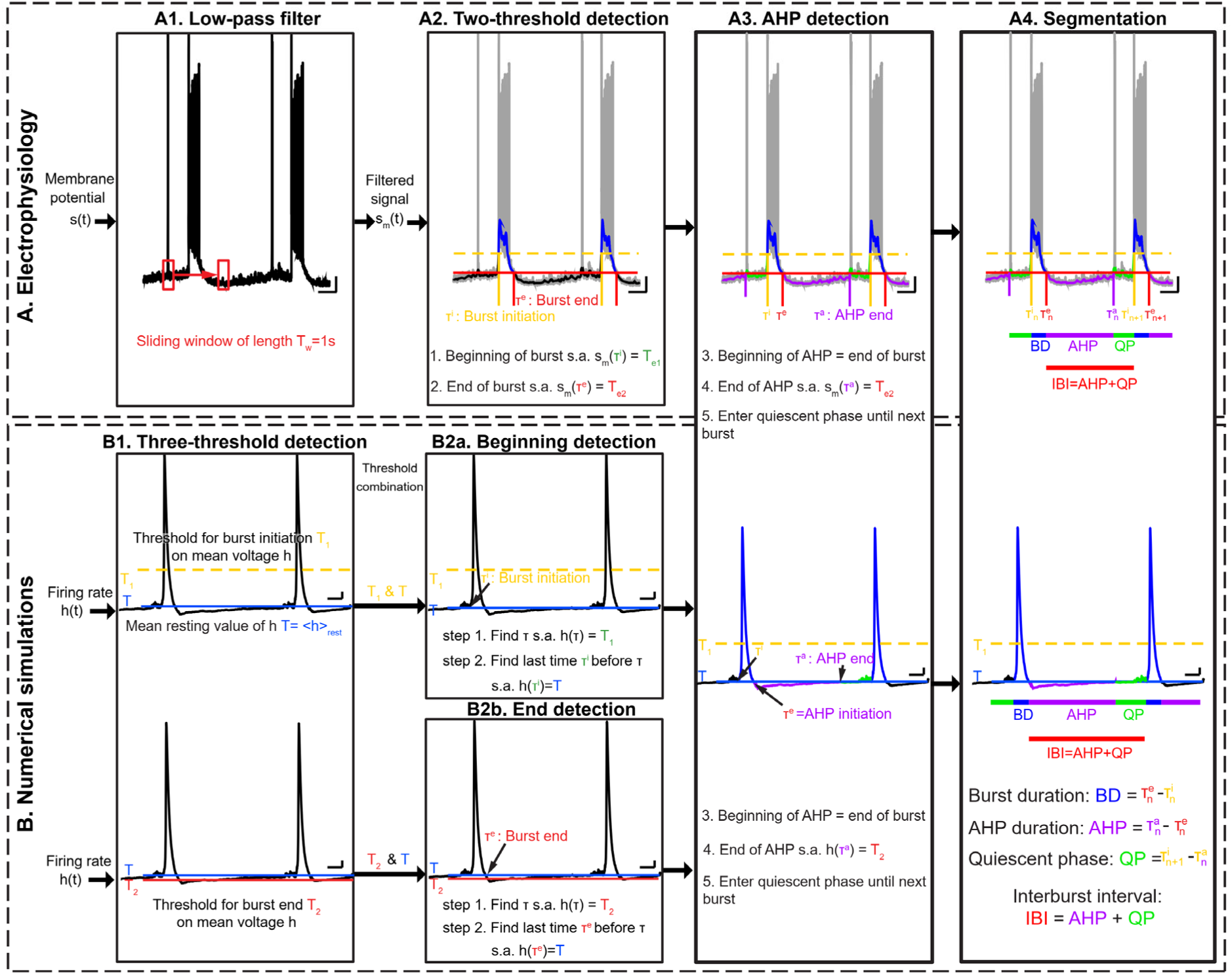
Burst detection algorithms. **A**) Burst detection for patch-clamp electrophysiological traces: the membrane potential signal is filtered using a sliding window of length *T_w_* = 1 s to apply a low-pass filter (A1), then a threshold detection is applied to the filtered signal *s_m_* (A2). AHP (3) begins at the end of the burst and lasts until *s_m_* is back above resting membrane potential, where the QP begins until the next burst. Extraction of burst durations (BD, blue), AHP duration (magenta) and QP (green) is then performed (4). IBIs are composed of AHP and QP. Scale bar: 10 s, 10 mV. **B**) Detection algorithm for simulated traces. Burst beginning is detected combining two thresholds, *T_1_* (yellow) and *T* (blue), on the mean voltage *h* (B1, top, and B2a), and burst end is detected combining a threshold *T_2_* (red) on facilitation *x* and threshold *T* on *h* (B1, bottom, and B2b). 3-4: AHP and QP are detected similarly as in the experimental traces, by segmenting the traces in three periods (BD, AHP and QP; 3-4). Scale bar: 10 s, 0.1 (normalized amplitude).

**Fig. S4.**
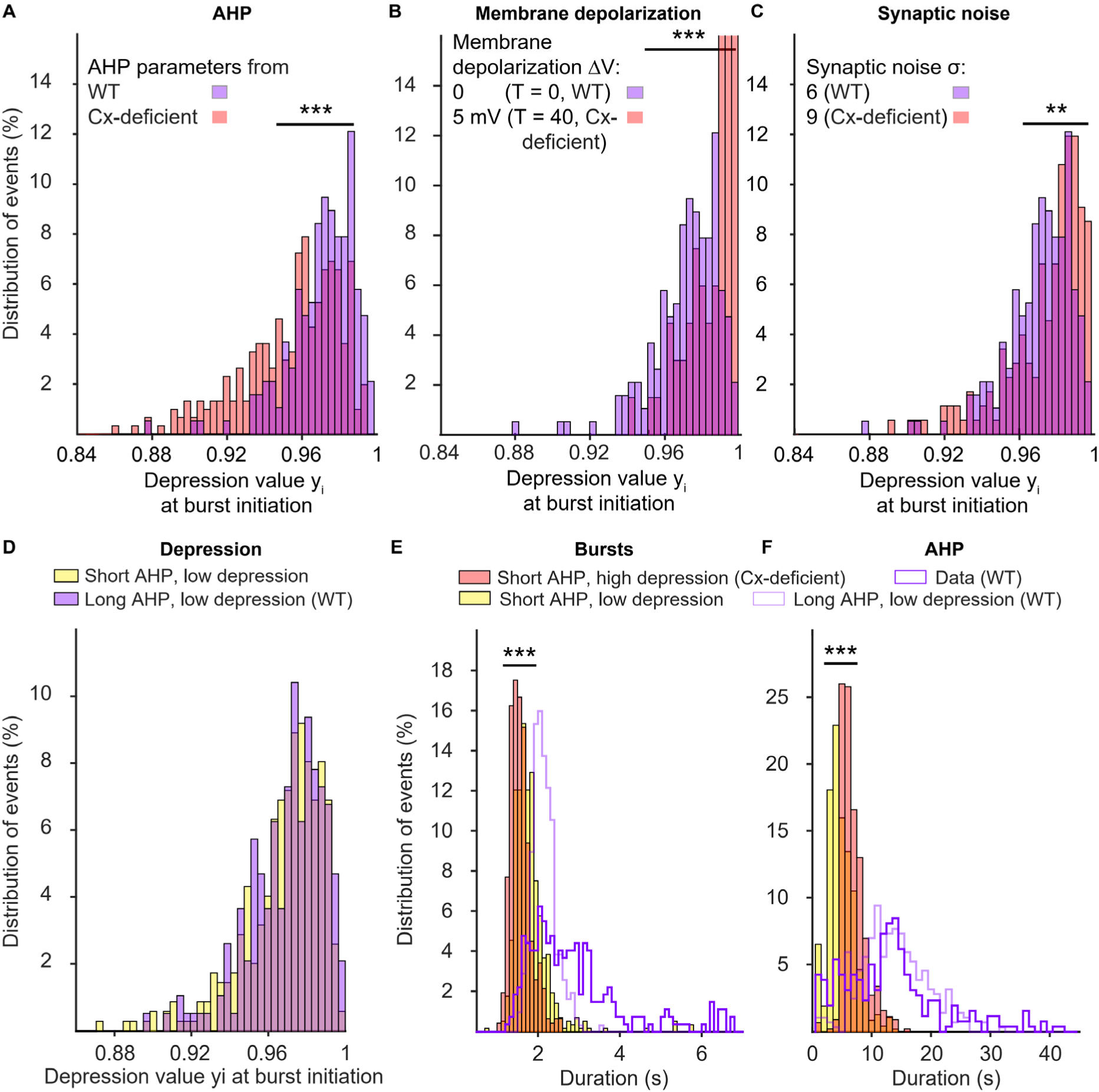
Synaptic depression: dependence on AHP, noise and depolarization and influence on burst dynamics. **A**) Distribution of depression level at burst initiation from the 5000 s simulations with τ_sAHP_ = 10.5 s, τ_mAHP_ = 0.35 s, T_AHP_ = − 30 (WT, light purple), and for τ_sAHP_ = 7.5 s, τ_mAHP_ = 0.15 s, T_AHP_ = − 23 (Cx-deficient, light red). σ = 6 and T = 0 for both conditions (p < 0.001, two-sample Kolmogorov-Smirnov test). **B**) Distribution of depression level at burst initiation from the 5000 s simulations with τ_sAHP_ = 10.5 s, τ_mAHP_ = 0.35 s, T_AHP_ = − 30, σ = 6 and T = 0 (WT, light purple) or T = 40 (Cx-deficient, light red) (p < 0.001, two-sample Kolmogorov-Smirnov test). **C**) Distribution of depression level at burst initiation from the 5000 s simulations with τ_sAHP_ = 10.5 s, τ_mAHP_ = 0.35 s, T_AHP_ = − 30, T = 0 and σ = 6 (WT, light purple) or σ = 9 (Cx-deficient, light red) (p = 0.0013, two-sample Kolmogorov-Smirnov test). **D)** Distribution of depression level at burst initiation from the 5000 s simulations with τ*_sAHP_* = 10.5 s, τ*_mAHP_* = 0.35 s, *T_AHP_* = − 30, *σ* = 6, *T* = 0 and with τ*_r_* = 2.9 s (light purple) and for τ*_sAHP_* = 5 s, τ*_mAHP_* = 0.15 s, *T_AHP_* = − 2 3, *σ* = 6, *T* = 0 and with τ*_r_* = 1.9 s (light yellow; p = 0.614, two-sample Kolmogorov-Smirnov test). **E**) Distribution of burst durations for τ*_sAHP_* = 5 s, τ*_mAHP_* = 0.15 s, *T_AHP_* = -23, *σ* = 6 and *T* = 0 (light red, Cx-deficient) and reduced depression (light yellow; p < 0.0001, Two-sample Kolmogorov-Smirnov test). The distribution of burst durations for WT parameters is indicated by the light purple line. **F)** Distribution of AHP, with the same parameters as in (**E**). The purple curves represent the distribution of burst (**E**) and AHP (**F**) durations obtained from experimental data in WT mice. Asterisks indicate statistical significance (***, p < 0.0001; **, p < 0.001).

